# Development of a highly efficient base editing system for *Lactobacilli* to improve probiotics and dissect essential functions

**DOI:** 10.1101/2024.08.12.607654

**Authors:** Hitoshi Mitsunobu, Yudai Kita, Yumiko Nambu-Nishida, Shoko Miyazaki, Kensuke Nakajima, Ken-ichiro Taoka, Akihiko Kondo, Keiji Nishida

## Abstract

*Lactobacilli* play essential roles in the food industry and are increasingly explored for their potential as probiotics and therapeutic agents. Beneficial strains are primarily isolated from various natural sources including healthy human bodies, and undergo rigorous characterization and safety evaluations. Genomic and genetic information has increasingly accumulated and been linked to their various functions, to which transgenic approaches are being performed to verify crucial genes. In order to reasonably develop more useful strains, beneficial traits need to be introduced into any given strains and enhanced or combined. However, for practical use as probiotics or foods, organisms with transgene are hardly acceptable. Here, we have introduced the base editing Target-AID system specifically for *Lactobacilli*, enabling precise installation of point mutations without donor DNA and at multiple genomic loci simultaneously. *Lactiplantibacillus plantarum* has been successfully engineered to reduce production of imidazole propionate, which has been reported to be associated with type 2 diabetes. Additionally, this system enabled transient knock-out of an essential gene, such as one involved in cell division showing severe filamentous cell phenotype, providing a unique approach for dissecting essential gene function.

**Importance:** This work provides highly efficient and multiplexable base editing system that installs precise point mutations in the genomes of the two major *Lactobacilli* strains. As the advanced CRISPR technology so-called non-cleaving genome editing, base editing is less toxic and does not integrate any foreign DNA into the genomes. Our approaches pave the way for dissecting and improving probiotics and food-grade microbes, ultimately creating better human health.

## Introduction

*Lactobacilli* are gram-positive, rod-shaped lactic acid bacteria and facultative anaerobes. They constitute an important part of the microbiome in various parts of the human body (1, 2) and are utilized in the production of fermented food products such as cheese and yogurt, which are recognized for their positive health effects. Due to their longstanding safe use in human consumption, *Lactobacilli* are generally recognized as safe (GRAS) microorganisms. They have been reported to play roles in improving gut microbiota, modulating immune function, and mitigating stress-induced phycological disorders, prompting their investigation as probiotics and clinical applications (3–7). Despite their beneficial effects, the genetic and molecular mechanisms underlying these health-promoting properties are yet to be fully understood. In order to utilize *Lactobacilli* as safe and more effective probiotics and therapeutics, rational engineering tools are much needed not only to elucidate the mechanism of action as well as to improve strains with enhanced efficacy (2).

CRISPR-Cas9 system has emerged as a powerful tool due to its efficiency and versatility in genetic manipulation (8, 9). Cas9 induces DNA double strand breaks (DSBs) at specific sites targeted by guide RNA (gRNA), which is then subjected to DNA repair process by the host cell. Once correctly repaired, the site is retargeted until an incorrect repair is made and a mutation is introduced. To facilitate incorporation of desired mutation, donor DNA can be co-introduced that contains homologous arms and intended mutation. However, in many bacterial species, the cytotoxicity associated with the nuclease activity of Cas9 and their poor DSB repair activities often limit its practical application (10). Multiplex editing is more challenging because DNA cleavage at multiple sites is synergistically more toxic, and donor DNA at each site is also required, making the process more complex and less effective. In *Lactobacilli*, nickase version of Cas9, which cleaves only single strand of DNA has been used to mitigate cytotoxic effects in combination with donor DNA (11, 12).

To circumvent the toxicity of DSB and the need for donor DNA, base editing technology has been developed by employing DNA deaminases tethered to nuclease-impaired CRISPR systems. Cytosine or adenine deaminase mediates base conversion within a narrow window in target sites, introducing specific point mutations (13, 14). Base editing has been applied not only eukaryotes but also in several laboratory bacterial species with robust performances (15–19). As base editing does not use donor DNA, it is easier to multiplex and the mutants may be classified as non-transgenic in many regions.

Here, we have applied and optimized base editing method for *Lactobacillus* species using the Target-AID system, a cytosine base editor that employs the cytosine deaminase from sea lamprey and uracil glycosylase inhibitor (UGI) (15, 20–22). The optimized systems achieved almost 100% editing efficiencies in *Lactiplantibacillus plantarum* and *Lactobacillus gasseri* strains and can be multiplexed. This allowed us to generate of a strain with improved property for human health, as well as to perform functional analysis of an essential gene by transient knock-out.

## Results and Discussion

### Development of base editing system in *Lactiplantibacillus plantarum*

To develop an efficient base-editing system in *Lactobacillus* species, we constructed a series of plasmids expressing various composition of essential components: nickase Cas9 (D10A: nCas), PmCDA1 deaminase, UGI, crRNA, and tracrRNA (Fig. 1A) with the replication origin from the *Lactobacillus-E. coli* shuttle vector pIB184 (23). The effector proteins (i.e. nCas, PmCDA1 and UGI) are expressed as a fusion protein or separate proteins by a single promoter in a monocistronic fashion. The crRNA that contains targeting sequence with scaffold was positioned downstream of the effector genes and is driven by a separate promoter. The tracrRNA was expressed either a bidirectional promoter shared with the effector proteins or a separate promoter. In some constructs terminators are added downstream of the intended transcripts. To evaluate the editing efficiency of the constructs, we first selected the uracil phosphoribosyltransferase (*upp*) gene as a target, a non-essential gene whose loss of function confers resistance to 5-fluorouracil in *L. plantarum* WCFS1. In the target sequence of 23 bases, cytosines at distal side of PAM were expected to be edited to thymine (Fig. 1B). The initial construct pYK01 employed the promoters previously used for expressing CRISPR-Cas9 system in *Lactobacillus* species (24–26) but achieved poor editing frequency after transformation and sequencing of 8 randomly selected colonies (Fig. 1C and 1D). Various composition of expression cassettes were then made and tested to find out that addition of a terminator at the end of crRNA yields nearly 100% editing efficiencies (Fig. 1D: pYK06‒08 and supplementary Fig. 1). The terminator might play a role in increasing half-life of transcripts and/or blocking unintended read-through to downstream sequences that interfere with plasmid function.

**Fig 1.**
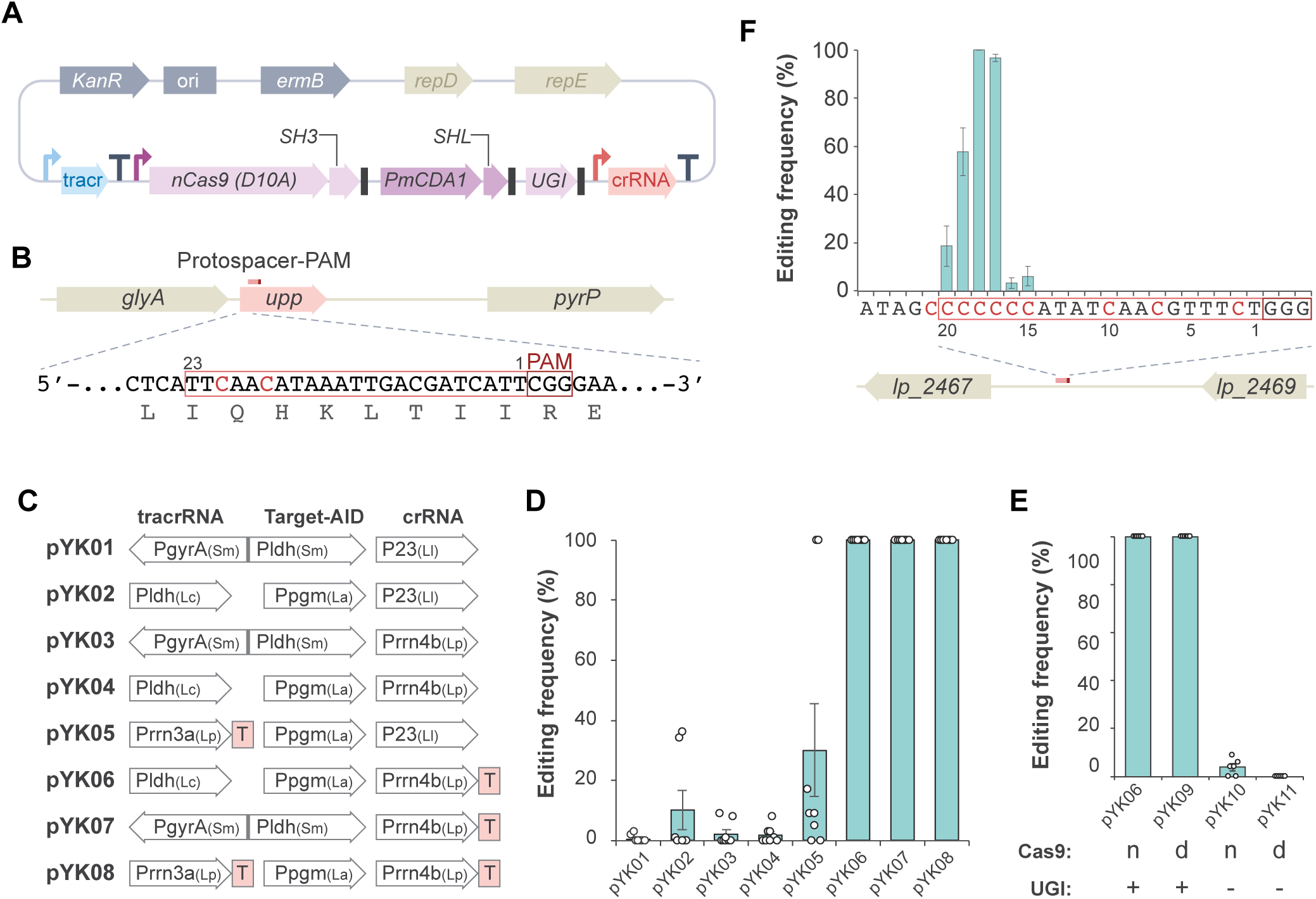
Development of Target-AID base editing system in *L. plantarum*. (A) Target-AID plasmid based on the *Lactobacillus-E. coli* shuttle vector pIB184. TracrRNA and crRNA are expressed as distinct transcripts as dual gRNA system. nCas9, PmCDA1 and UGI are expressed as a single transcript and are translated into separate proteins, as stop codons (black boxes) are placed at the ends of each coding sequence. To mediate the binding of nCas9 and PmCDA1, the SH3 domain and its ligand, SHL, are fused to the respective proteins. The angled arrows represent the promoters. Terminators (T) are placed as necessary. (B) The target sequence design for *upp* gene. The 23 bases of protospacer sequence with PAM are boxed. Cytosines that are subjected to base editing are shown in red. Amino acid sequences are shown at the bottom. (C) The Target-AID plasmid variants with different combination of promoters and terminators. The derived organism for promoters is described in the parentheses. Sm, Ll, Lc, La, and Lp stand for *Streptococcus mutans*, *Lactococcus lactis*, *Lacticaseibacillus paracasei*, *Lactobacillus acidophilus*, and *Lactiplantibacillus plantarum*, respectively. (D) The editing frequencies of each plasmid are measured by sequencing of eight randomly selected colonies and plotted. The bars and error bars represent the mean and standard deviation, respectively. Independent spectra data for pYK02 and pYK06 are shown in Supplementary Fig. 1. (E) Effect of nCas9, dCas9, and UGI on editing frequency. (F) Editing window of Target-AID in *L. plantarum*. A 20 bases target sequence (boxed) containing multiple cytosines (red) was selected from the intergenic region and edited by using pYK6. Averaged editing frequencies (bars) at each base position are shown with standard deviation (error bars).

In base editing systems, nCas9 is generally preferred to dCas9 as it is more efficient. This is because nicking the opposite strand of a deaminated base inhibits the base excision repair of the deaminated base, facilitating insertion of the mutated base. On the other hand, nCas9 shows somewhat toxicity, especially in bacteria, which can be inferred from the reduced transformation efficiency. To see if nCas9 is necessary for efficient base editing in *Lactobacilli*, dCas9 version was tested showing comparable efficiency to nCas9 version as long as it contains UGI (Fig. 1E), which inhibits the initial step of base excision repair. Thus, the dCas9 version can be used in cases such as using a strain exhibiting low transformation efficiency.

### Determination of editing window in *L. plantarum*

Each base editing system has a unique editing window, meaning that efficient base conversions occur within a limited range of bases within the target sequence. Editing window for Target-AID has been shown to be at approximately 16–20 bases away from a PAM in other organisms (20, 21, 27) . To determine the editing window of pYK06 in *L. plantarum*, we selected a target site containing multiple cytosines at positions 15–21 from a PAM at intergenic region (Fig. 1F). Consistently, base conversion activity was prominently observed at positions 17–20, with the most efficient editing occurring at cytosines in positions 17 and 18. Several studies in other organisms also have shown that different editing windows and base-change patterns can be selected by using different base-editing systems.

### Expanding targeting scope with NG-Cas9 in *L. plantarum*

PAM sequence requirement limits the selection of targetable sequence. The original CRISPR-Cas9 system from *Streptococcus pyogenes* requires a PAM sequence of NGG at the 3’end of target sequences. Engineering studies on the Cas9 protein have produced variants with altered PAM requirements, among which NG-Cas9 requires reasonably simplified NG while maintaining substantial editing activity (28). By adopting the variant, we intended to develop the Target-AID-NG plasmid for *Lactobacilli*. However, during construction of the requisite plasmid in *E. coli*, the spacer sequence in the crRNA, which encodes target sequence, was found to be frequently mutated and appeared to be self-edited. Due to this effect, we could only obtain a mixed population of plasmids containing the correct spacer sequence and mutated. Although the spacer sequence in the plasmid is followed by GA at its 3’ end and not supposed to be recognized by NG-Cas9, the engineered variant might have ambiguity in the recognition and have self-edited (Supplementary Fig. 2A). Using such mixed population of plasmids, we transformed *L. plantarum* to assess its editing capacity at target sites harboring either NGG or NG PAMs (Supplementary Fig. 2B). For the Cas9-NG Target-AID system (pYK12, mixed population), we could show editing at the both target sites, albeit with lower efficiency compared to the canonical Target-AID for NGG site (Supplementary Fig. 2C), presumably due to impurity of the plasmid. Similar self-targeting effect has been documented for base editing in *E.coli* (29) and plants (30), which could be circumvented by using alternative gRNA scaffold sequence starting GCCCC (30). Nonetheless, despite these challenges, Target-AID-NG enabled the acquisition of edited clones with practical usability in *Lactobacillus*.

### Efficient multiplex base editing in *L. plantarum*

Multiplex editing using nuclease in bacteria is challenging because it causes severe toxicity and requires donor DNA for each site. As base editing is less toxic and can be multiplexed by multiple crRNA only, we have designed plasmids containing single, dual, or triple crRNA cassettes introduced tandemly in the plasmids (pYK06-s, -d, or -t; Fig. 2A, 2B and Supplementary Fig. 3). These constructs were transformed in *L. plantarum* with comparable transformation efficiencies and achieved highly efficient base conversions at the all of the targeted sites corresponding to the crRNA cassettes (Fig. 2C and supplementary Fig. 3). This system employs an original dual gRNA consisting of tracrRNA and crRNA, rather than a single-stranded chimeric guide RNA (sgRNA), which means that only the crRNA part is needed for multiplexing, and the shorter repeat sequence length facilitates plasmid construction. In addition, multiple crRNAs can be provided in a manner of CRISPR array, where the spacer sequences are flanked by the crRNA repeat sequences and transcribed under a single promoter regulation, which further decreases the size and complexity of the plasmid.

**Fig 2.**
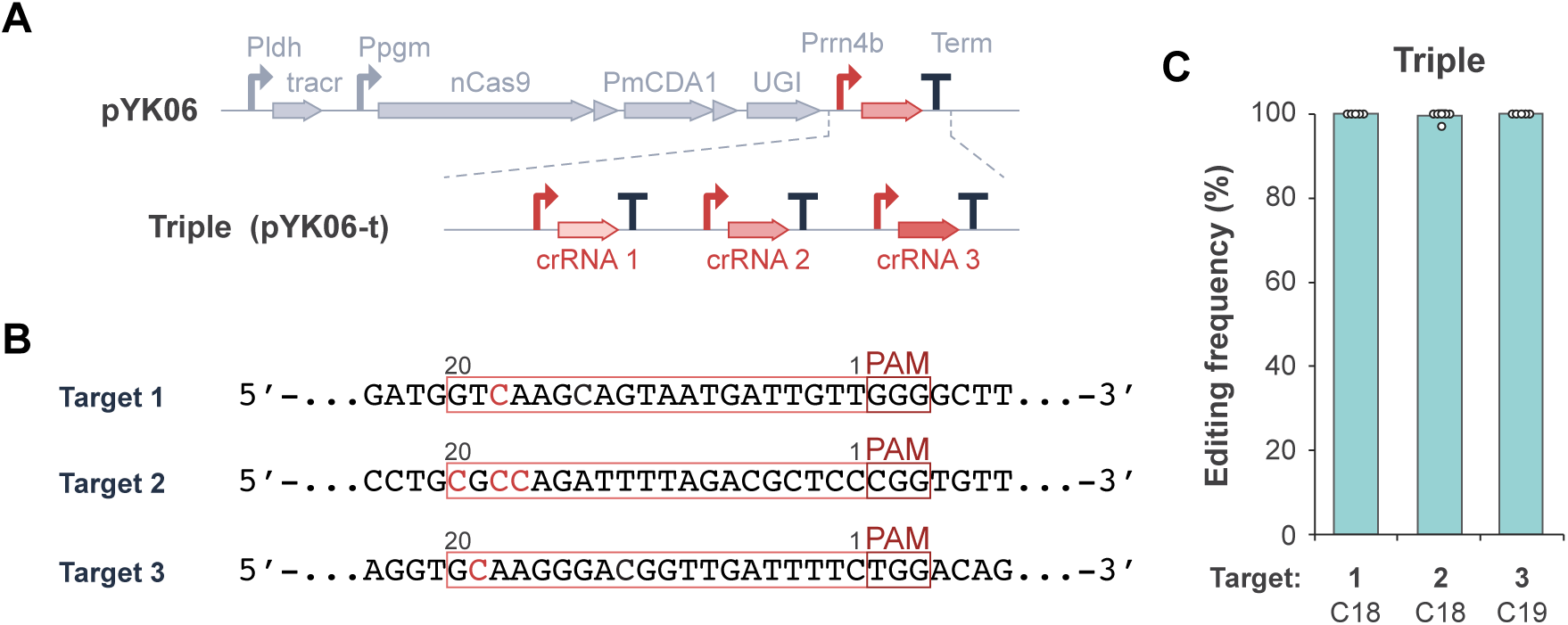
Multiplex Target-AID base editing in *L. plantarum.* (A) Each crRNA expression cassette contains a promoter, a crRNA with a target sequence and a terminator, inserted in tandem in pYK06. (B) Three target sequences are selected from *urdA* gene and editable cytosines are shown in red. (C) The editing frequencies at the three sites by pYK06-t are measured by sequencing of six randomly selected colonies and the value from the highest base positions were plotted. Averaged editing frequencies (bars) are shown with standard deviation (error bars). Full dataset is shown in Supplementary Fig. 3.

### Compatibility of the base editing system in *Lactobacillus gasseri*

As a wide variety of *Lactobacilli* are practically used, we sought to determine whether our base editing system would be compatible with other species and tested in *L. gasseri* ATCC 33323 strain, a human commensal strain found in various areas including oral and intestinal microbiota. A target sequence was designed to modify an intergenic region to circumvent unexpected effects on cell growth and/or transformation efficiency (Fig. 3A). While the transformation efficiency in *L. gasseri* is lower compared to *L. plantarum* under the conditions tested, we could obtain transformants with the pYK06 plasmid. However, none of the *L. gasseri* transformants showed detectable base conversion at the target site (0/8: Fig. 3C), suggesting that the regulatory elements in pYK06 plasmid may not work well in *L. gasseri*. Instead, pYK07 which contained PgyrA-Pldh bidirectional promoter performed well in *L. gasseri* (8/8 transformants: Fig. 3B, 3C, and Supplementary Fig. 4) as well as *L. plantarum* (Fig. 1D). This marked improvement underscores the importance of promoter selection in achieving successful genome editing outcomes across species.

**Fig 3.**
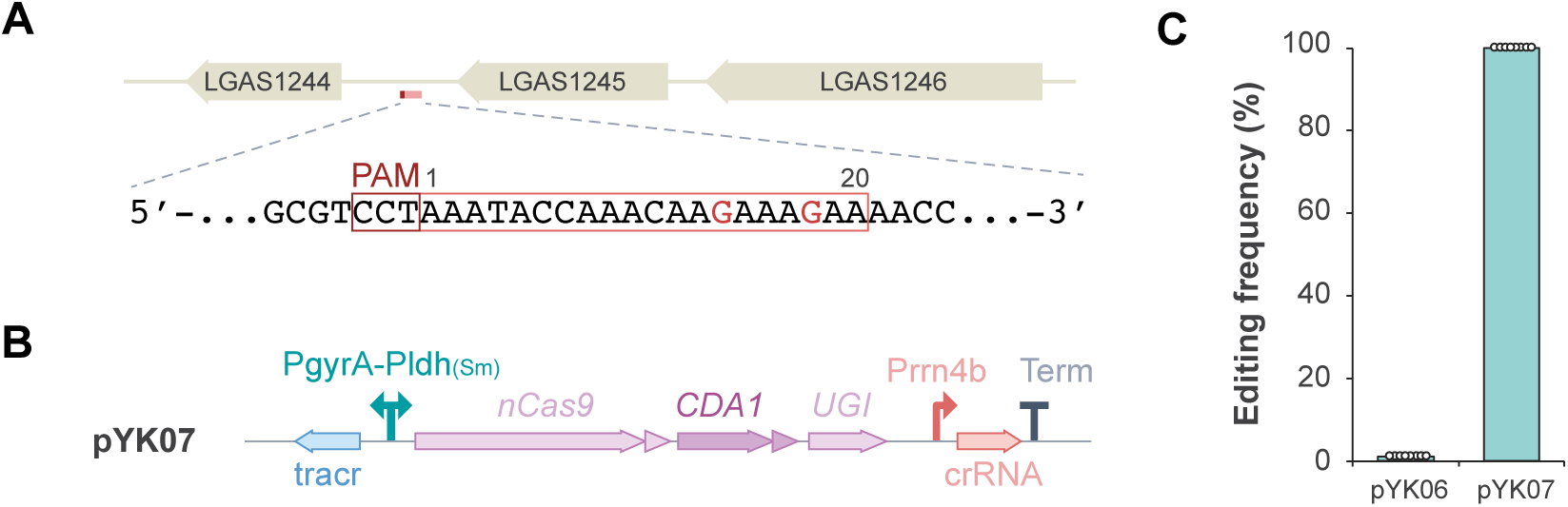
Target-AID base editing in *L. gasseri*. (A)A 20-bases target sequence was selected at intergenic region and the opposite strand is shown with the guanines subjected to editing and shown in red. (B) The architecture of Target-AID in pYK07. The PgyrA-Pldh _(Sm)_ promoter has bidirectional transcriptional activity and drives tracrRNA and effector protein transcript. (C) The editing frequencies of pYK6 and pYK7 in *L. gasseri*. are measured by sequencing of eight randomly selected colonies and plotted. Averaged editing frequencies (bars) at each base position are shown.

### Engineered *L. plantarum* strain with reduced imidazole propionate production

Many of *Lactobacilli* harbor *urdA* gene encoding urocanate reductase that produces imidazole propionate (ImP) from urocanate in histidine utilization pathway (31). It has been shown that microbially produced ImP is associated with type 2 diabetes, impairing glucose tolerance and insulin signaling (31, 32). To develop a *Lactobacillus* strain that does not produce ImP and can be used for probiotics and foods, we designed crRNA for *urdA* to introduce a premature stop codon in *L. plantarum* using pYK06 (Fig. 4A). After transformation, almost 100% editing efficiency was confirmed in all four randomly selected clones (Supplementary Fig. 4), suggesting that the disruption of *urdA* did not cause severe growth defect in the condition tested. To establish transgene-free and stable strain, cells were grown in antibiotic-free medium until they lost antibiotic resistance associated with the plasmid. Lack of the plasmid was confirmed by PCR, and the mutation at the gene was also confirmed (data not shown). To determine the ability to produce ImP, cells were grown to saturation and the supernatant was obtained, which was subjected to liquid chromatography coupled to tandem mass spectrometry, showing more than 10-fold reduction in ImP production in the edited strains compared to wild-type (Fig. 4B). This indicates that *Lactobacilli* can be manipulated to reduce ImP production without impairing their growth, and could be used in foods and probiotics that may help control type II diabetes. Without integration of foreign DNA sequences, genome-edited organisms basically resemble those obtained by random mutagenesis (33), and can be considered as non-genetically modified organism in several regions including Japan and US, whereas the EU-court so far does not allow edited organisms by genome editing methods (including CRISPR-Cas) as ‘non-GMO’ (Callaway, 2018; Spök et al., 2022; Waltz, 2016).

**Fig 4.**
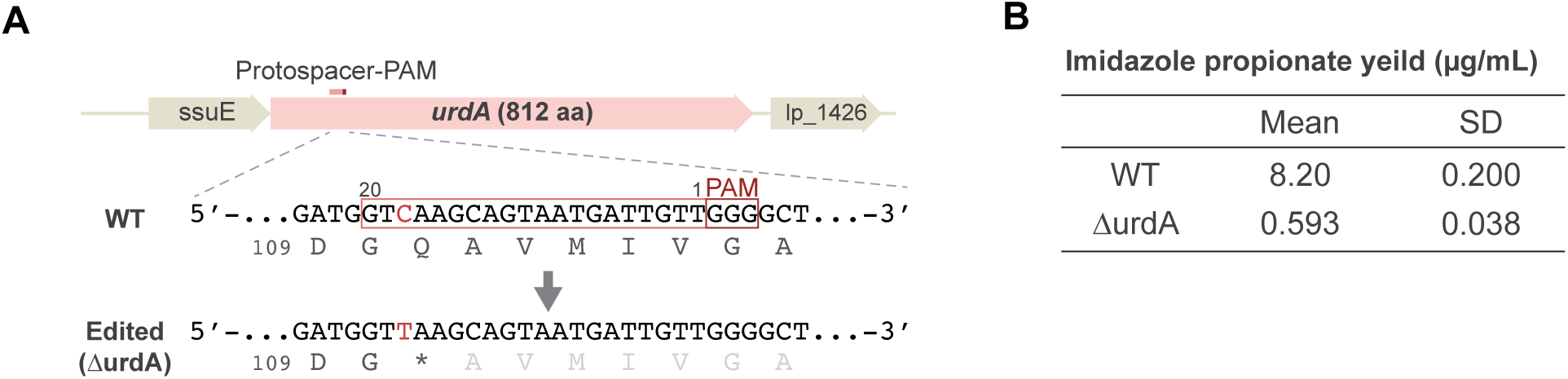
Establishment of urdA-edited *L. plantarum* strain with reduced imidazole propionate (ImP) production. (A) Target sequence was selected to install a stop codon at 111th glutamine (Q) in *urdA* gene. (B) The production of ImP by wild type (WT) and *urdA*-edited strain. Yields were measured in the growth media as described in Materials and Methods.

### Transient editing and analysis of an essential gene

The morphology of commensal microbes has been reported to be important for their colonization and stability in the gut ecosystem (37), but their genetic background has not been investigated outside of model bacteria. FtsZ is a master regulator of bacterial cell division and its conditional knockout mutants have been shown to have an elongated filamentous cell phenotype. Using pYK06, we designed seven gRNAs targeting the *ftsZ* in *L. plantarum* to introduce amino acid substitutions or stop codons at various positions in the FtsZ coding sequence (Fig. 5A), as its complete loss of function could be lethal. Intended edits were detected in target 2, 3, 4, 5, and 7 by sequencing of the colonies of obtained transformants (Fig. 5B). These mutants exhibited distinct filamentous morphological changes under optical microscopy (Fig. 5C), indicating that the *ftsZ* function was disturbed. We have tried to obtain a stable mutant devoid of the plasmid, but the recovered plasmid-free cells no longer contained any of the mutations. This suggests that these mutations were lethal in a long term in *L. plantarum* and the cells were in mixed population of edited and wild type cells, that were being edited in real-time, allowing transient monitoring of their mutant phenotype. Given that no mutations were found in targets 1 and 6, it is possible that these mutations are more severe and cause dominant-negative effects and cells may not survive even for a short period of time. This can be an indicator that a mutation is lethal if it is not obtained or is only obtained in such a transient manner and can be a unique methodology of transient generation and functional analysis of mutants with lethality or defect in growth to dissect essential cellular functions.

**Fig 5.**
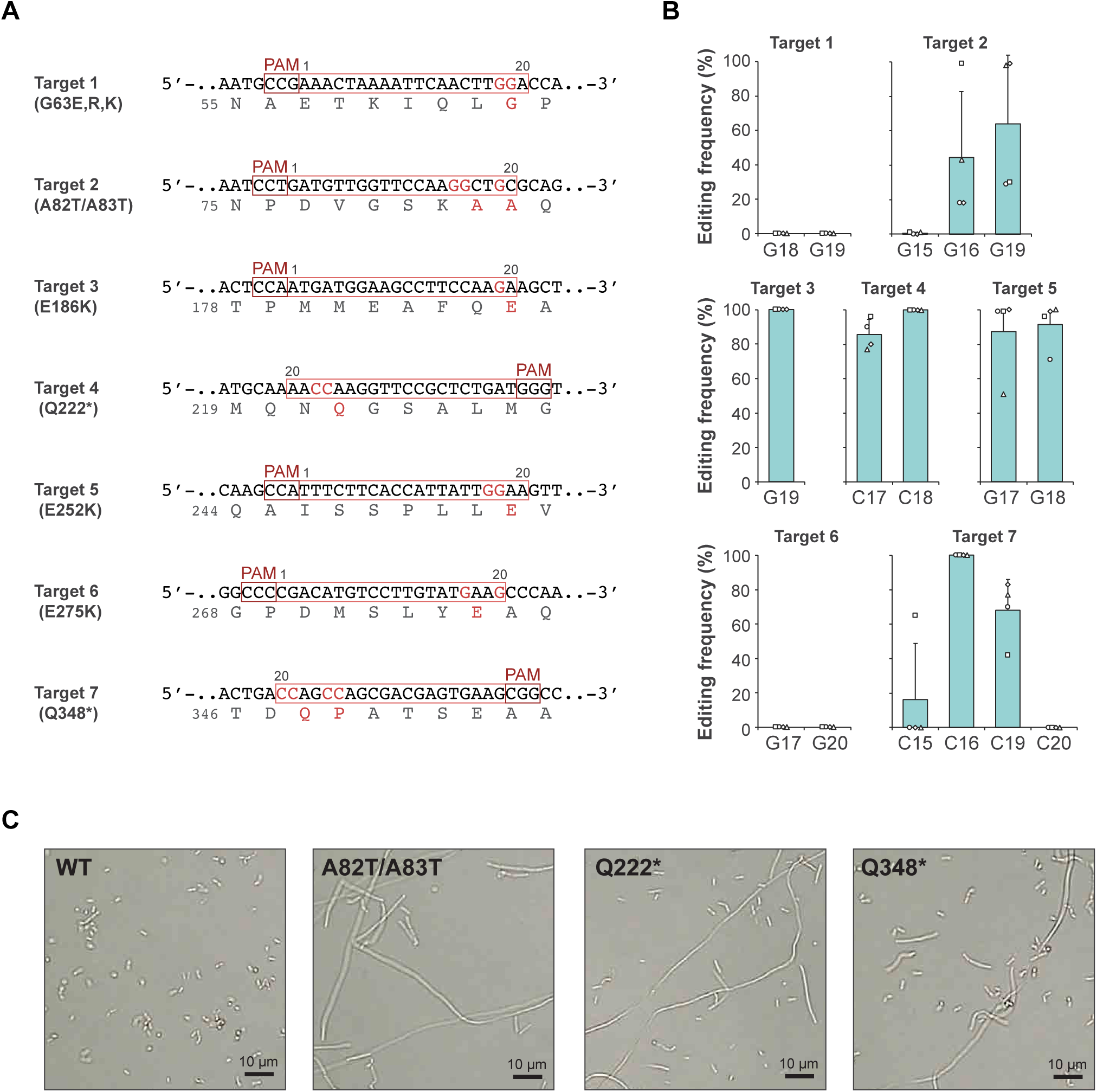
Editing of an essential cell division gene *ftsZ*. (A) Seven target sequences are designed to install amino acid substitutions or premature stop codons in FtsZ. (B) The editing frequencies at editable base positions for each target sequence was measured by sequencing four randomly selected colonies and plotted. Averaged editing frequencies (bars) at each base position are shown. (C) Optical microscopic images of *wild-type* and representative *ftsZ*-edited cells.

## Materials and methods

### Bacterial strains, plasmids, and media

Table 1 lists all bacterial strains and plasmids used in this study. The molecular cloning was carried out using conventional DNA ligation or Gibson Assembly (38). The promoters and the pIB184-derived region (*repD* and *repE* for replication in *Lactobacilli* and erythromycin resistant gene *ermB*) were synthesized by GenScript. PrimerSTAR Max polymerase (TaKaRa) and Tks Gflex polymerase (TaKaRa) were used for PCR amplification. Plasmid DNA was extracted using FastGene Plasmid Mini Kit (NIPPON Genetics Co, Ltd.). Restriction enzymes were purchased from New England Biolabs. Ligation high Ver.2 (TOYOBO) was used for ligation reactions. *E. coli* HST08 Premium Competent Cells (TaKaRa) were used for DNA vector amplification and grown in Luria-Bertani (LB) medium at 37°C with shaking. DNA sequences were confirmed by Sanger sequencing. The complete sequence of pYK06 and the different modules on the other plasmid are shown on Supplementary Note. *L. plantarum* WCFS1 and *L. gasseri* ATCC 33323 were cultured in de Man, Rogosa, and Sharpe (MRS) broth statically or MRS agar at 37°C. The erythromycin (Em) concentrations were 10 µg/ml for *Lactobacilli* and the kanamycin (Km) concentrations were 50 µg/ml for *E. coli*.

**Table 1.**
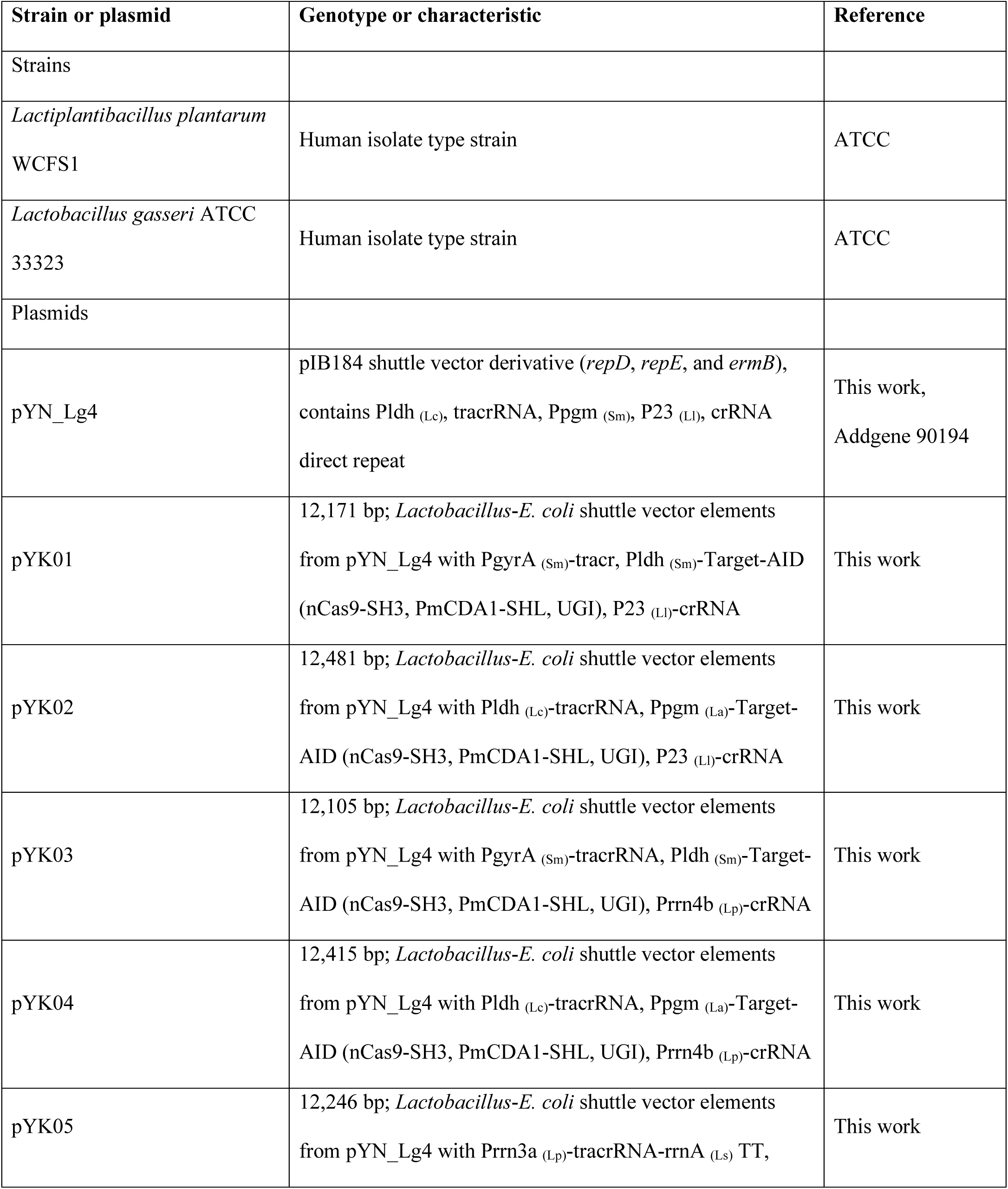
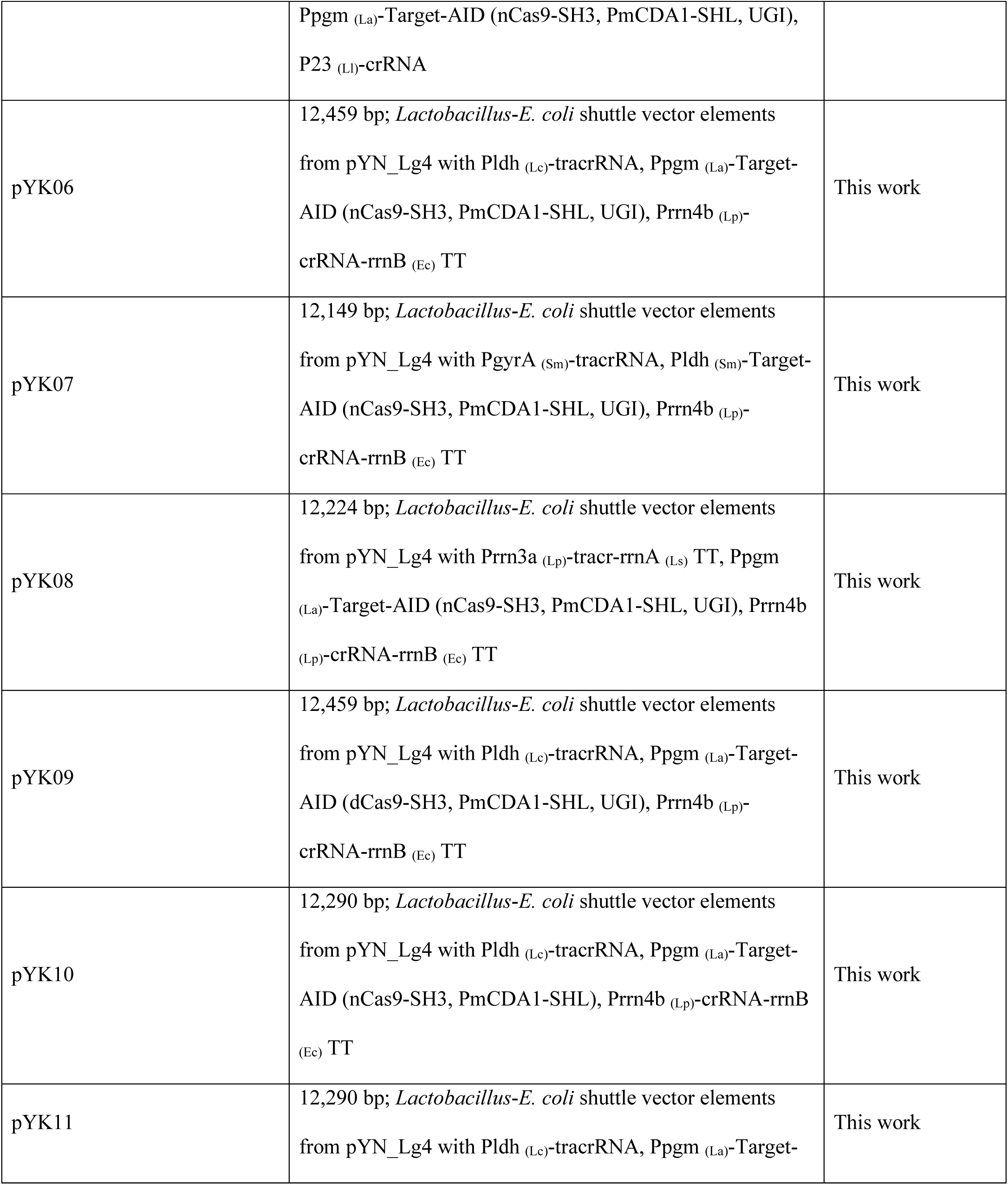
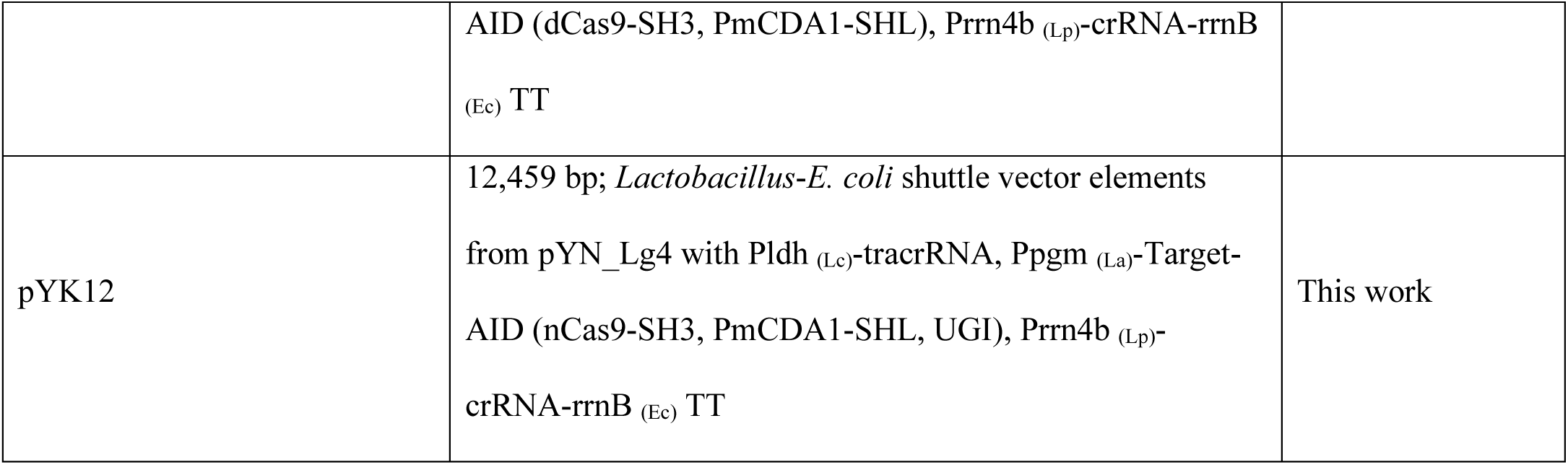
Bacterial strains and plasmids used in this study.

### Transformation of *Lactobacillus* strains

Transformation of the plasmids by electroporation of *L. plantarum* and *L. gasseri* was conducted by following the previous study (24). Two ml of the medium was inoculated with a single colony from a plate and incubated statically at 37°C overnight. One ml of this culture was back-diluted into 25 ml of MRS containing 0.41 M glycine in a 50 ml tube and was incubated statically at 37 °C until the OD_600_ reached approximately 0.85 (approximately 3.5 hours). Cells were centrifuged at 2,500 × g for 10 min at 4°C to pellet, washed twice with 5 ml of 10 mM MgCl_2_ followed by one wash with 5 ml of SacGly (10% glycerol with 0.5 M sucrose). The cells were then resuspended in 1 ml of SacGly and centrifuged at 20,000 × g for 1 min. After removing the supernatant, the final pellet was resuspended in 500 µl of SacGly. For electroporation, 60 µl of this suspension and plasmid (2–5 µg) were added to a 1 mm gap cuvette and subjected to electroporation using an ELEPO21 (NEPA GENE) with the following parameters: for poring pulse, voltage, 1250V; pulse width 2.5 msec; pulse interval, 50 msec; pulse frequency, 1 time; polarity, + and for transfer pulse: voltage, 150V; pulse width, 50 msec; pulse interval, 50 msec; pulse frequency, 5 times; polarity, +/-. After electroporation, 1 ml of MRS broth was added to the cuvette, transferred the cell suspension to a sterile tube, and incubated statically for recovery overnight at 30 °C. Subsequently, 250 µl of the recovered culture was plated on MRS agar supplemented with erythromycin.

### Mutational analysis

Isolated colonies from the MRS-agar plate supplemented with erythromycin were inoculated in 500 µl of MRS medium supplemented with erythromycin and allowed to grow at 30℃ overnight. The overnight culture was then spread on MRS-agar plate containing Em. A genomic region encompassing the target sites was PCR-amplified using EmeraldAmp PCR Master Mix (TaKaRa) with appropriate primers listed on Table 2 from randomly selected colonies and the resulting fragments were subjected to Sanger sequencing. The frequency of C-to-T change at the targets from each clone was calculated using EditR, an online base-editing analysis tool (39).

**Table 2.**
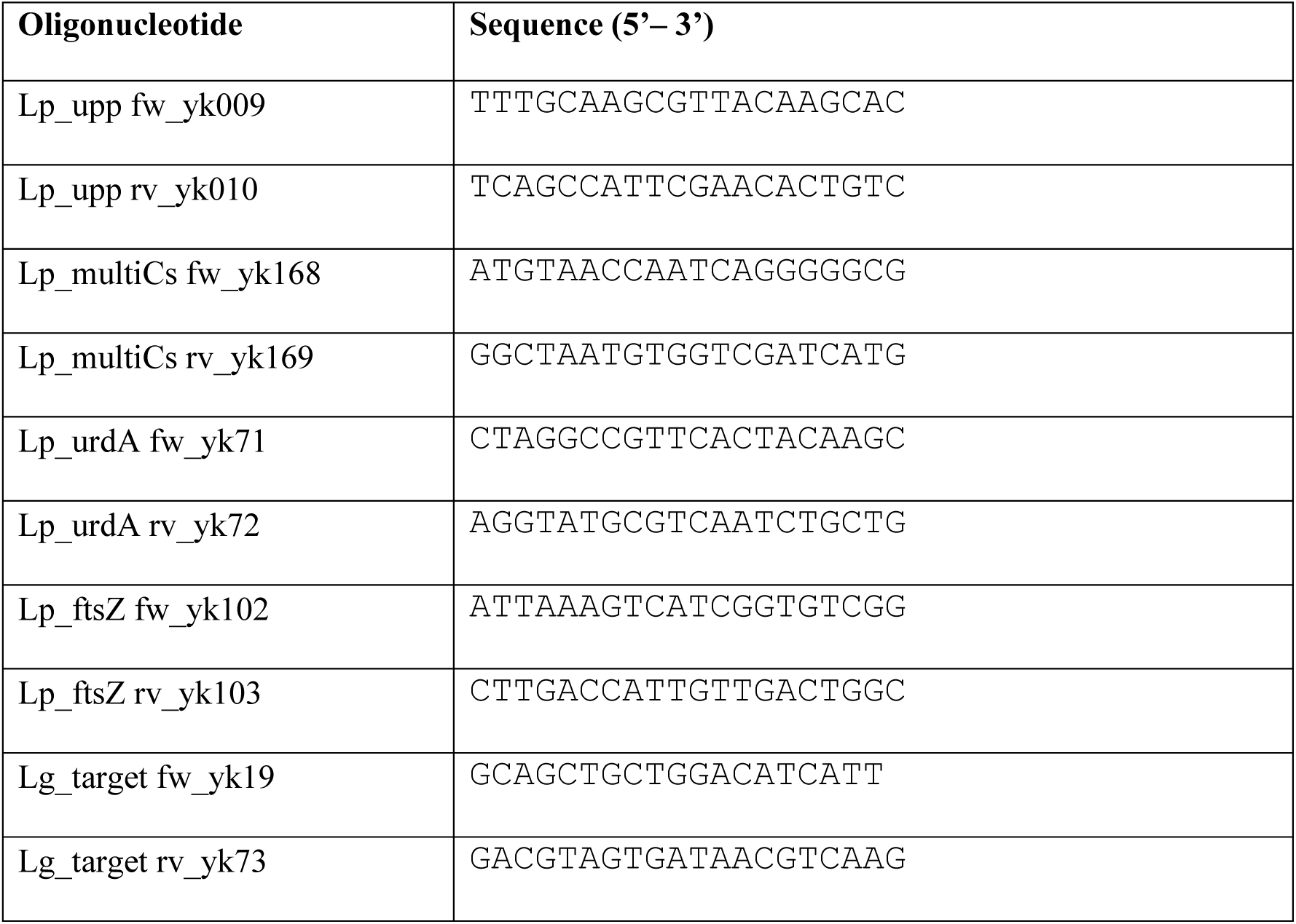
Oligonucleotides used in this study.

The highest editing frequency was adopted if multiple editable bases present at a target site. The editing frequency was determined as the average frequency of C-to-T changes across four to eight independent biological replicates.

### Microscopy of FtsZ mutants

To assess the cell morphology of the FtsZ mutants, a colony harboring the mutations was cultured in 500 µl of erythromycin-containing MRS media and incubated overnight at 30°C. Cell imaging was performed using a KEYENCE BZ-8000 microscope with a 40× objective lens.

### Evaluation of ImP production

To measure ImP production, 3 mL of Brain Heart Infusion (BHI) broth was inoculated with an isolated single colony of WT or the urdA-edited strain on MRS agar plates, and incubated statically at 37°C for approximately 15 hours until the OD600 reached 1.3–1.4 under the anaerobic condition. Thirty microliters of the 15-hr culture was added into 15 mL of BHI supplemented with 10 mM trans-urocanic acid. After incubation at 37°C statically for 24 hours under the anaerobic condition, the supernatant of the cell suspension was collected by centrifugation at 3,260 × g for 10 min. Five hundred microliters of each sample was mixed with 100 µL of water/acetonitrile/formic acid mixture (900:100:1) and 500 µL of acetonitrile, and applied to InterSep NH2 cartridge and Oasis PRiME HLB cartridge and the eluent was prepared in 10 mL of water/acetonitrile/formic acid mixture (900:100:1) as samples for LC-MS/MS analysis. LC–MS/MS analyses were performed on the Shimadzu UFLC system (CBM-20A/LC-20AD/SIL-30AC; Kyoto, Japan) with an InterSustain AQ-C18 150 × 2.1 mm, 5 µm column (GL science), coupled to a 4000QTRAP triple quadrupole mass spectrometer (SCIEX, Framingham, MA), with Turbo Ion Spray source with ESI ionization (electrospray ionization) in positive mode. Data were acquired by the software LabSolutions Version 5.118 (Shimadzu).

## Acknowledgments

This work was supported by Program on Open Innovation Platform with Enterprises, Research Institute and Academia (OPERA) Grant Number JPMJOP1851 to K.N., GteXProgram Japan Grant Number JPMJGX23B4 to K.N., Green Innovation funds of the New Energy and Industrial Technology Development Organization (NEDO) to K.N., the Japan Agency for Medical Research and Development (AMED) under Grant Number 21ek0109448h0002 and 24bk0104169s0201 to K.N.

## Supplementary Note 1

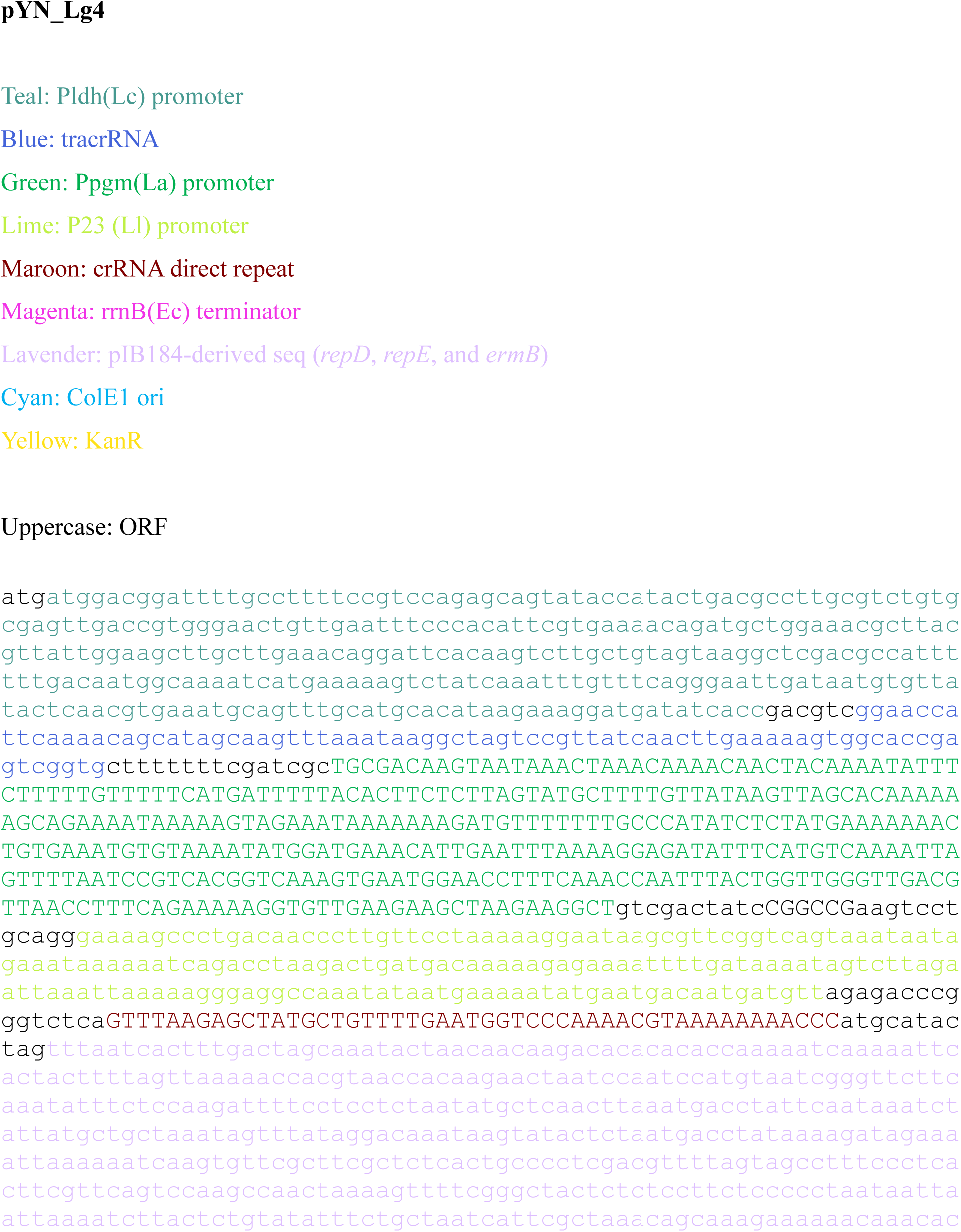
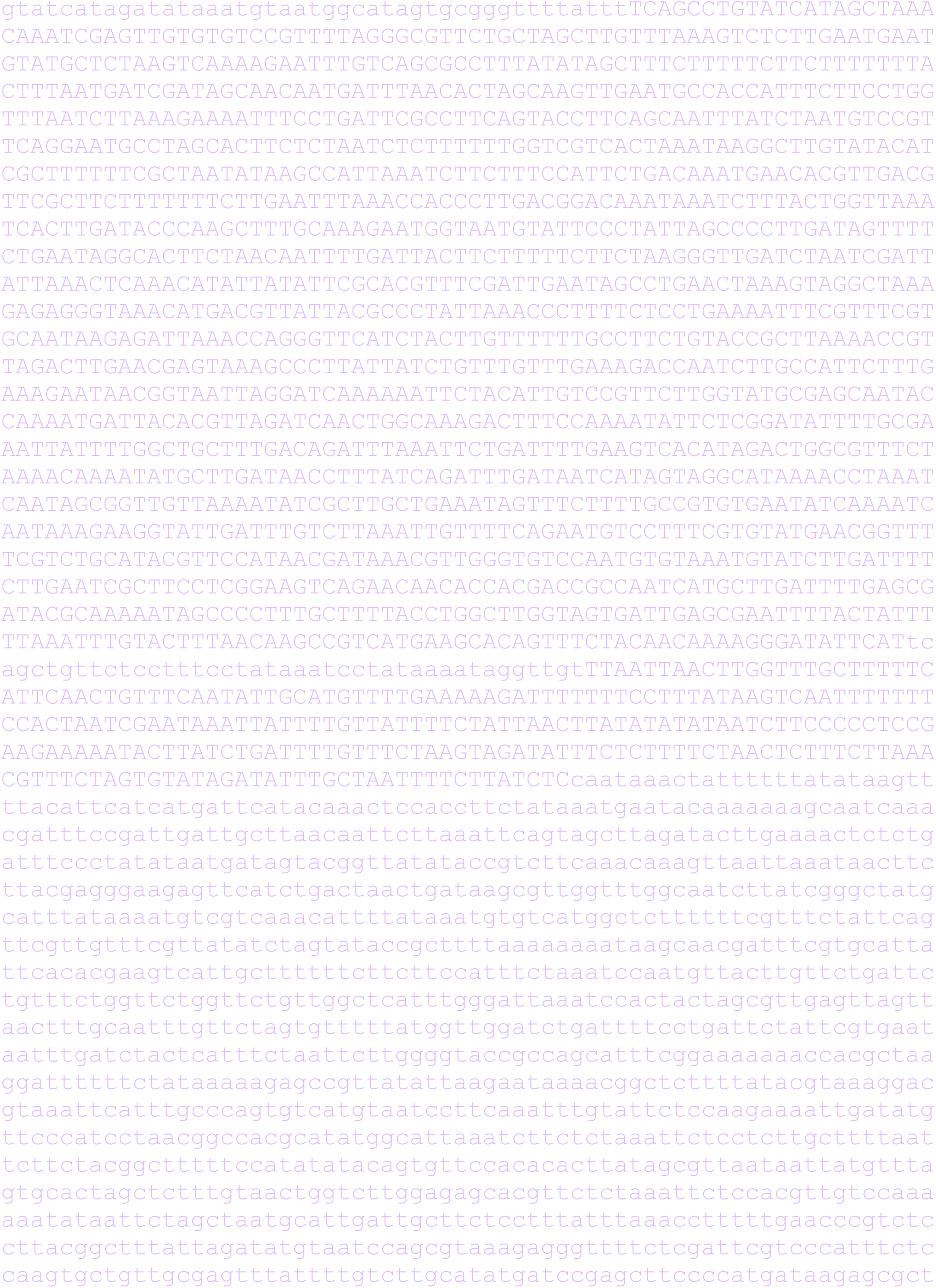
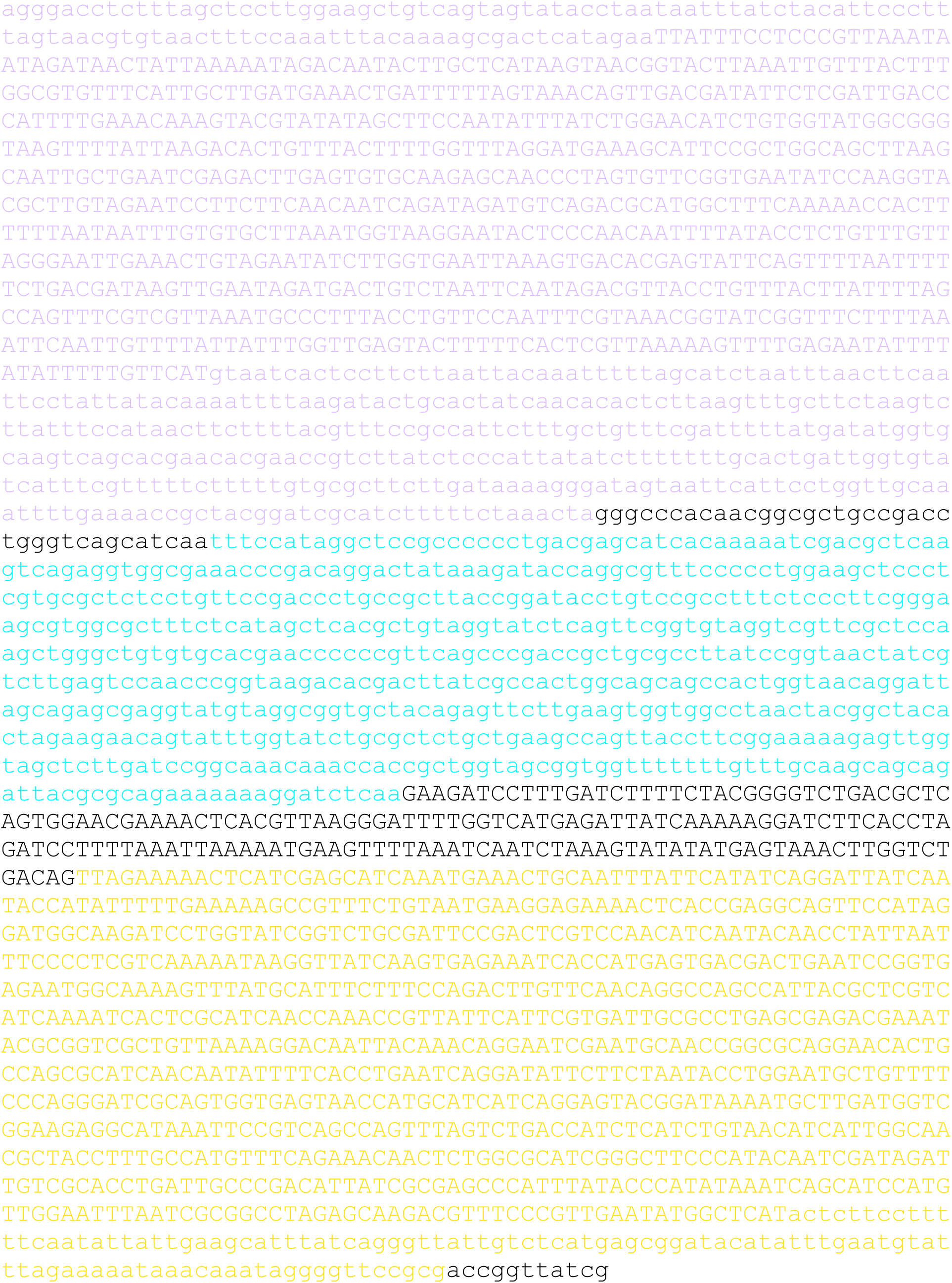

## Supplementary Note 2

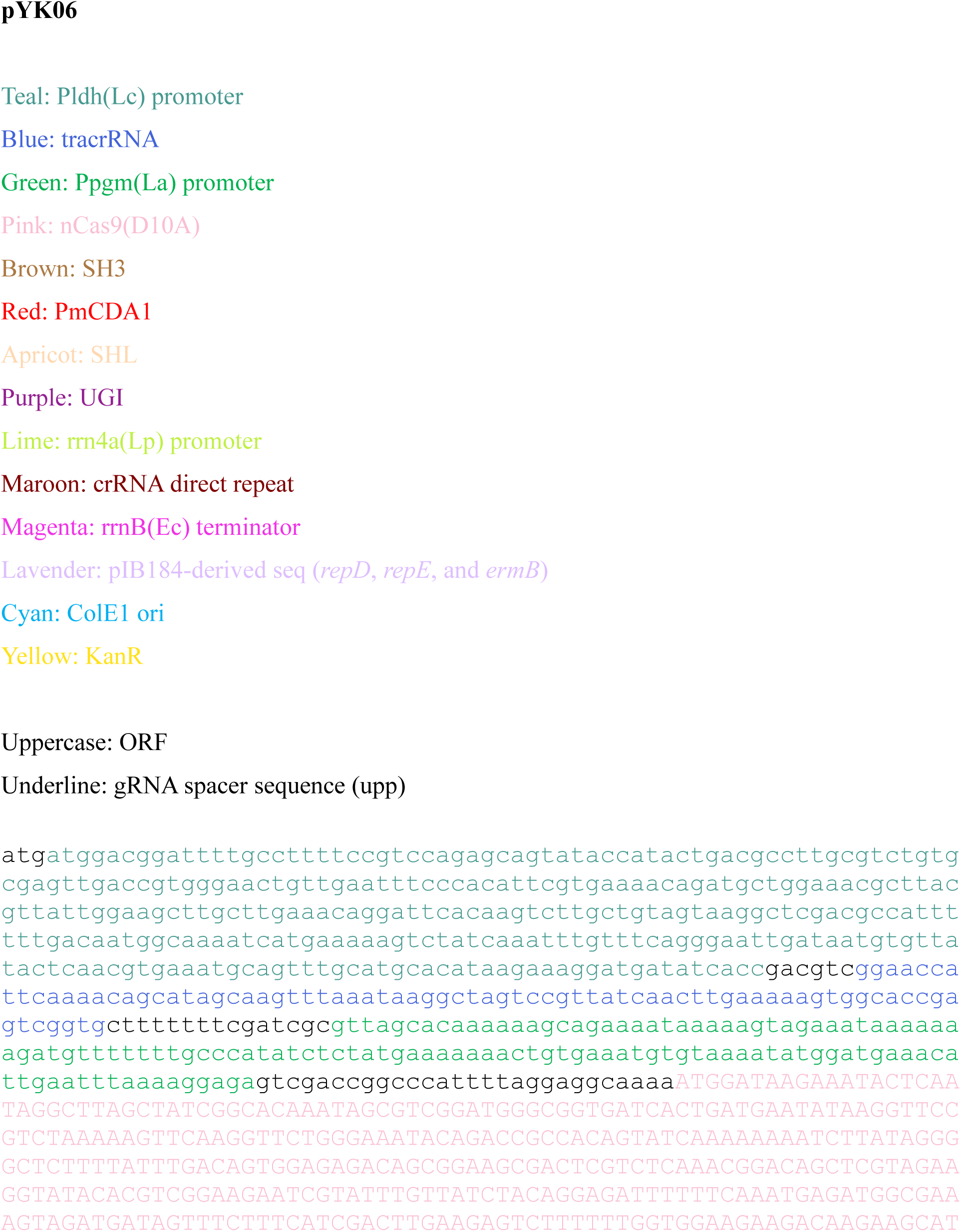
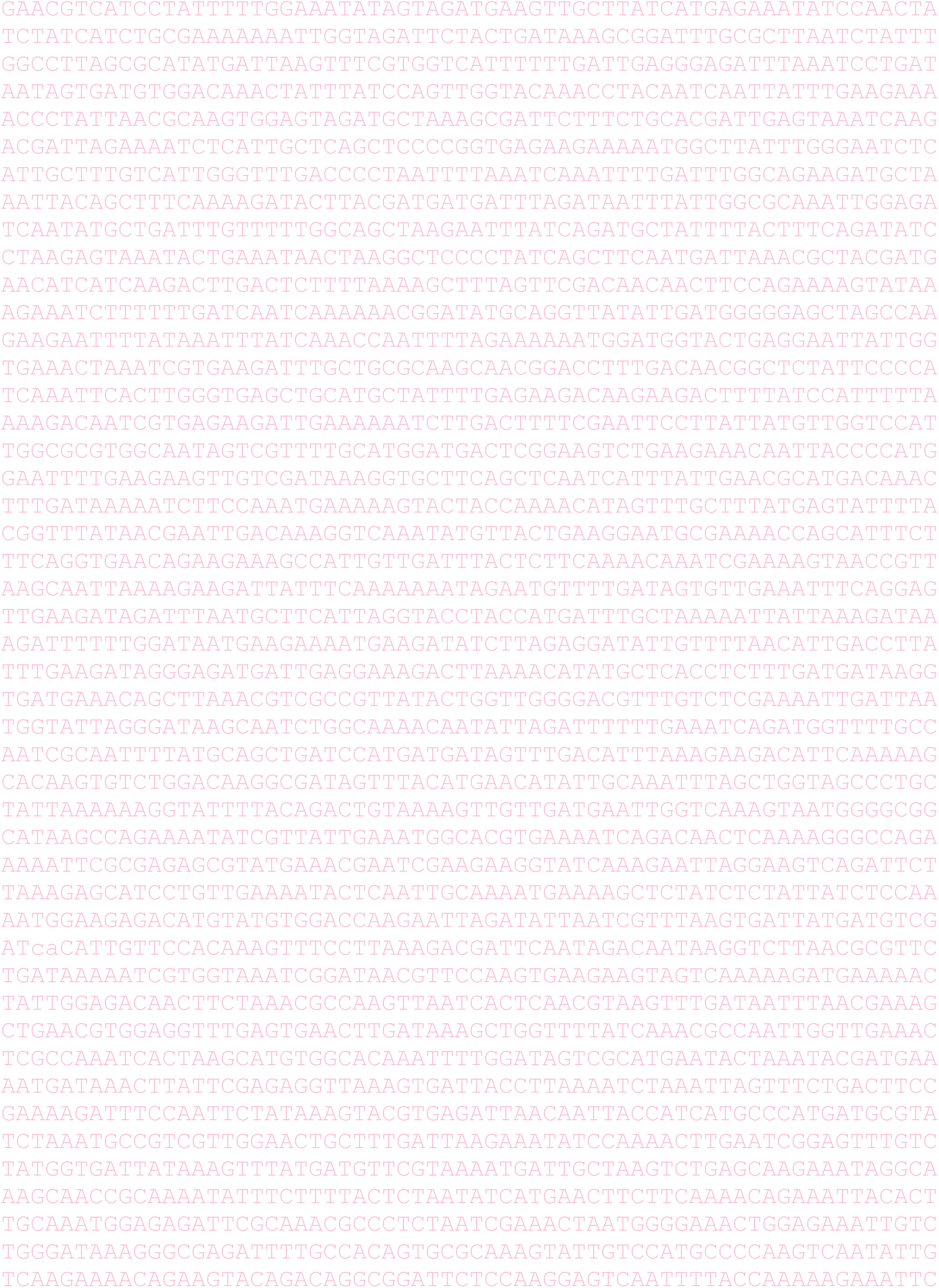
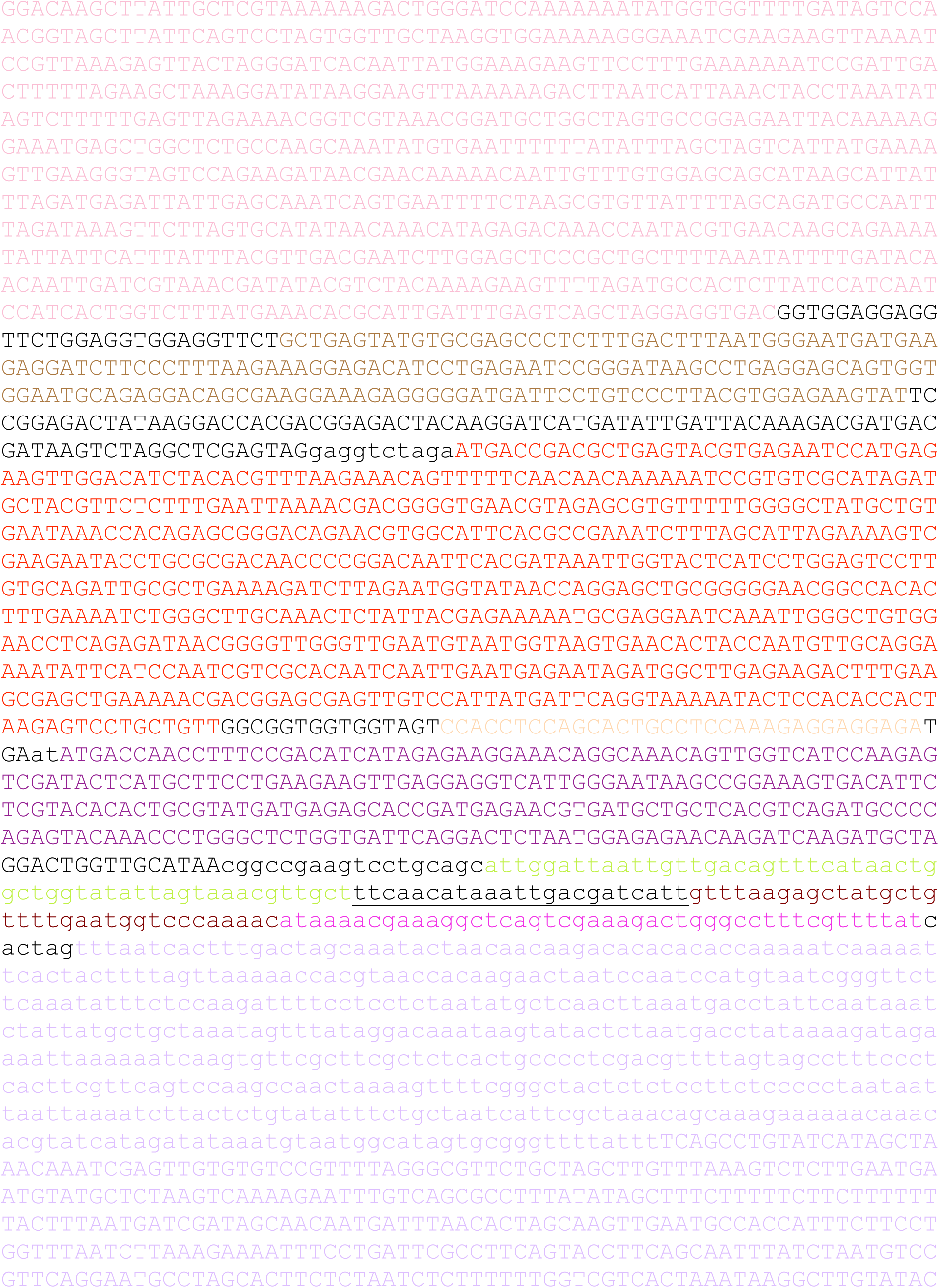
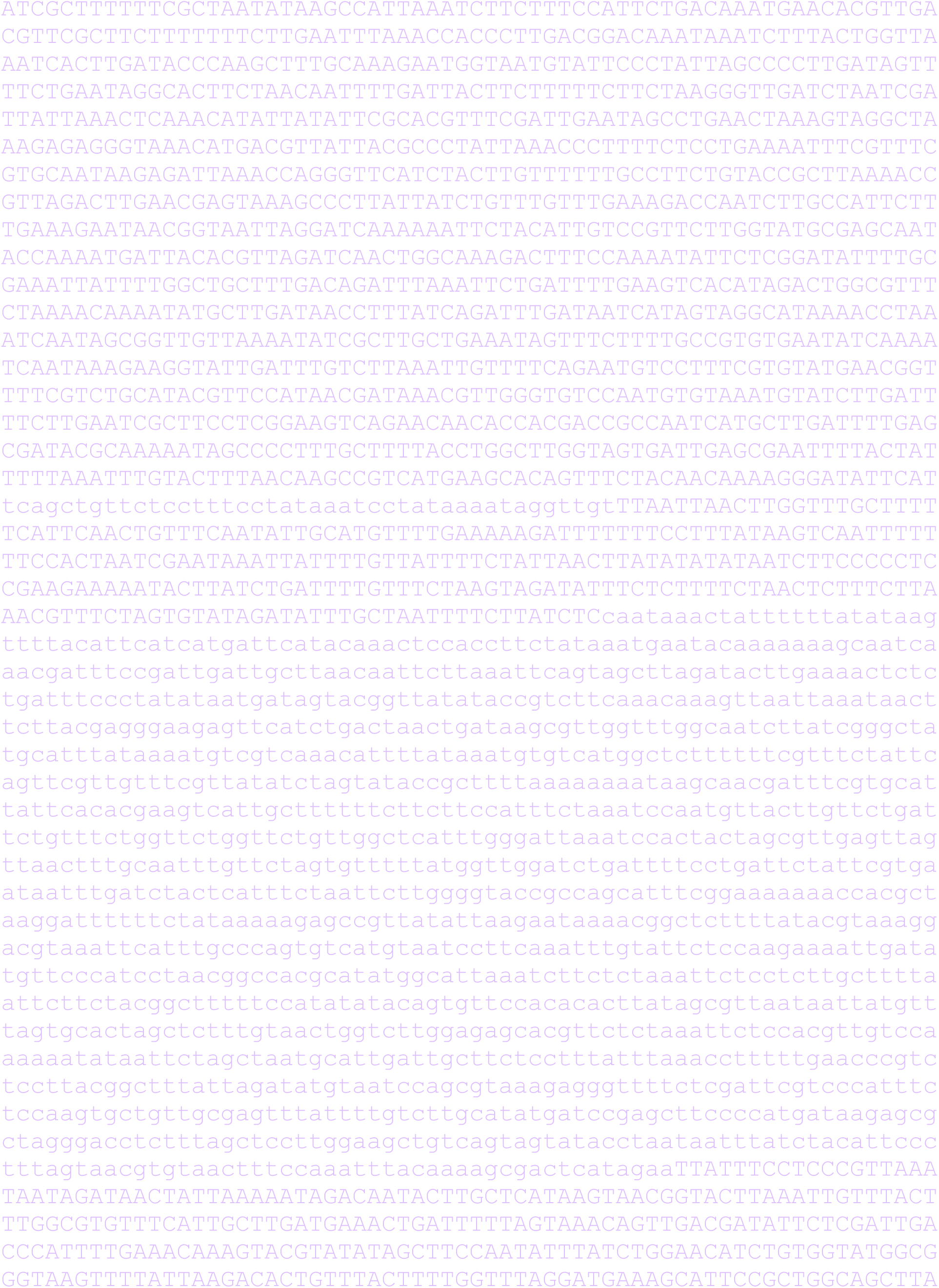
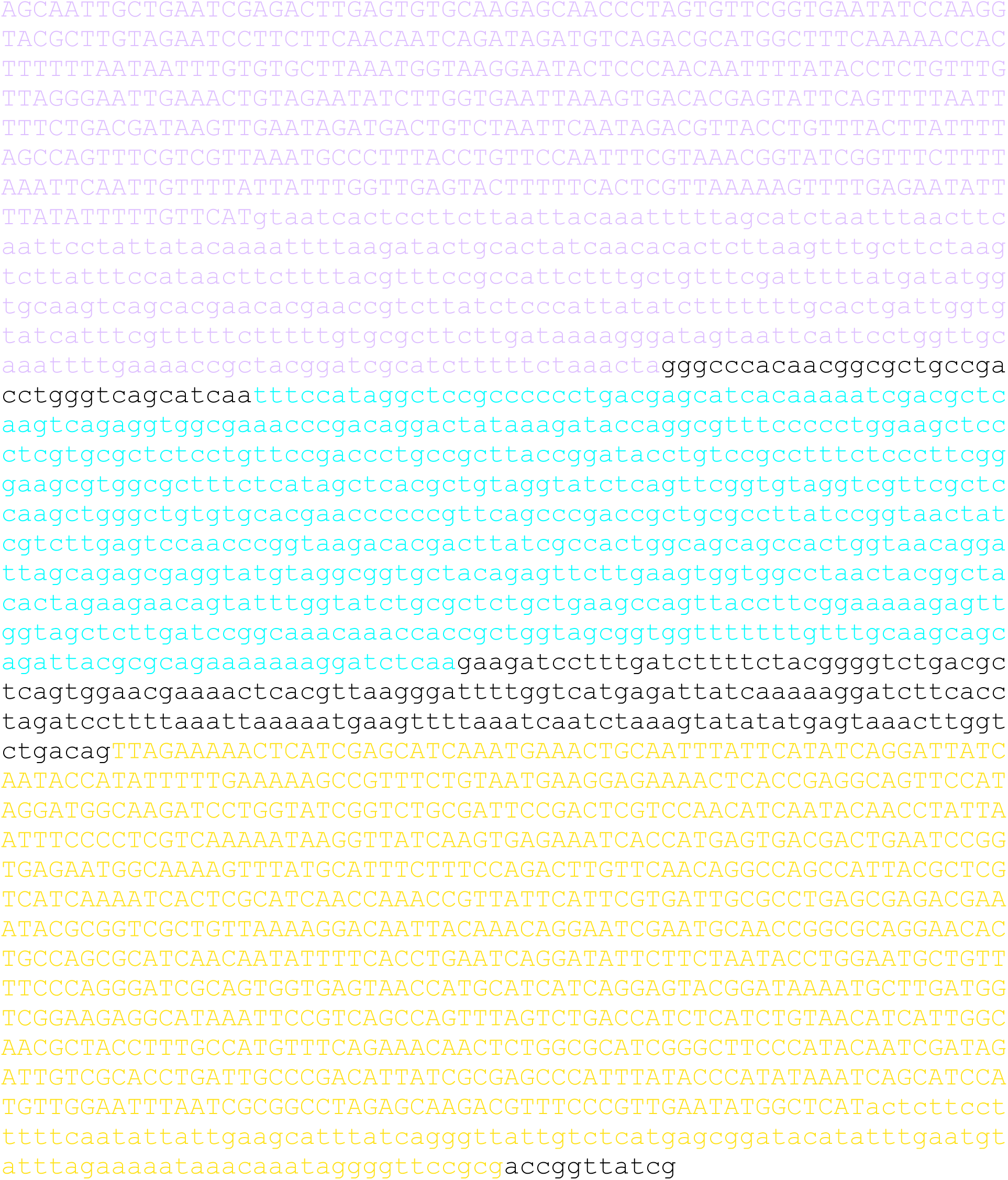

## Supplementary Note 3

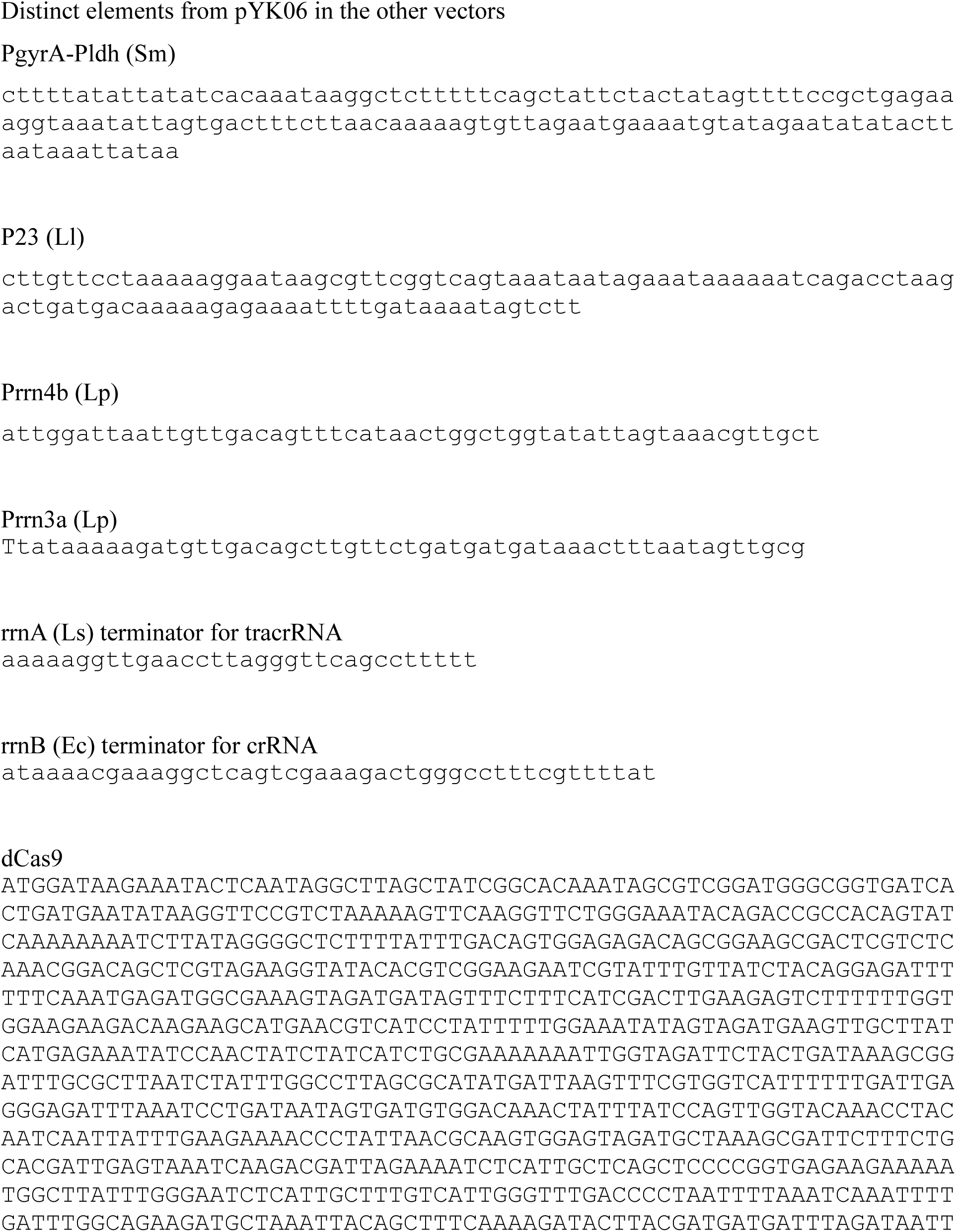
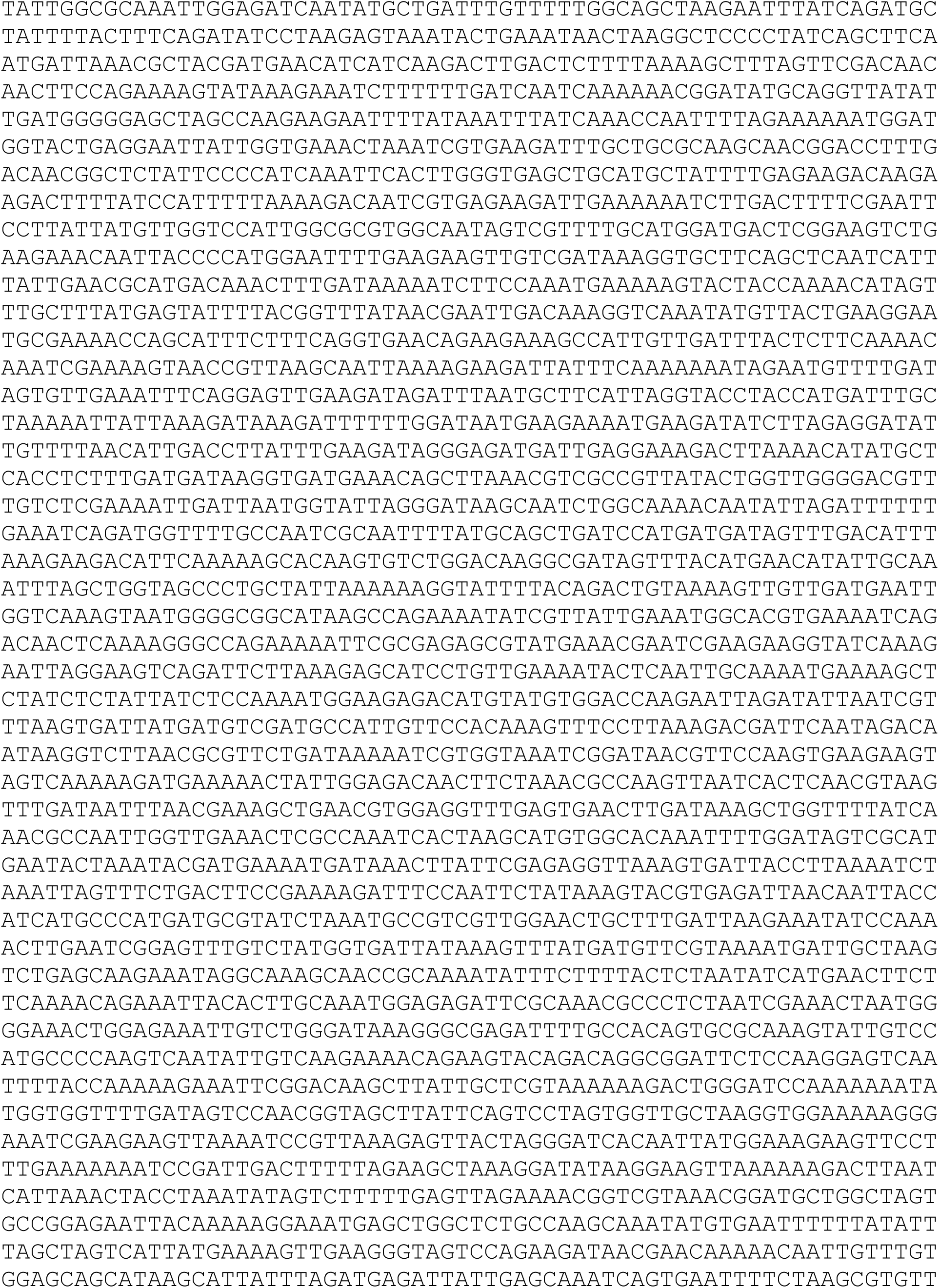
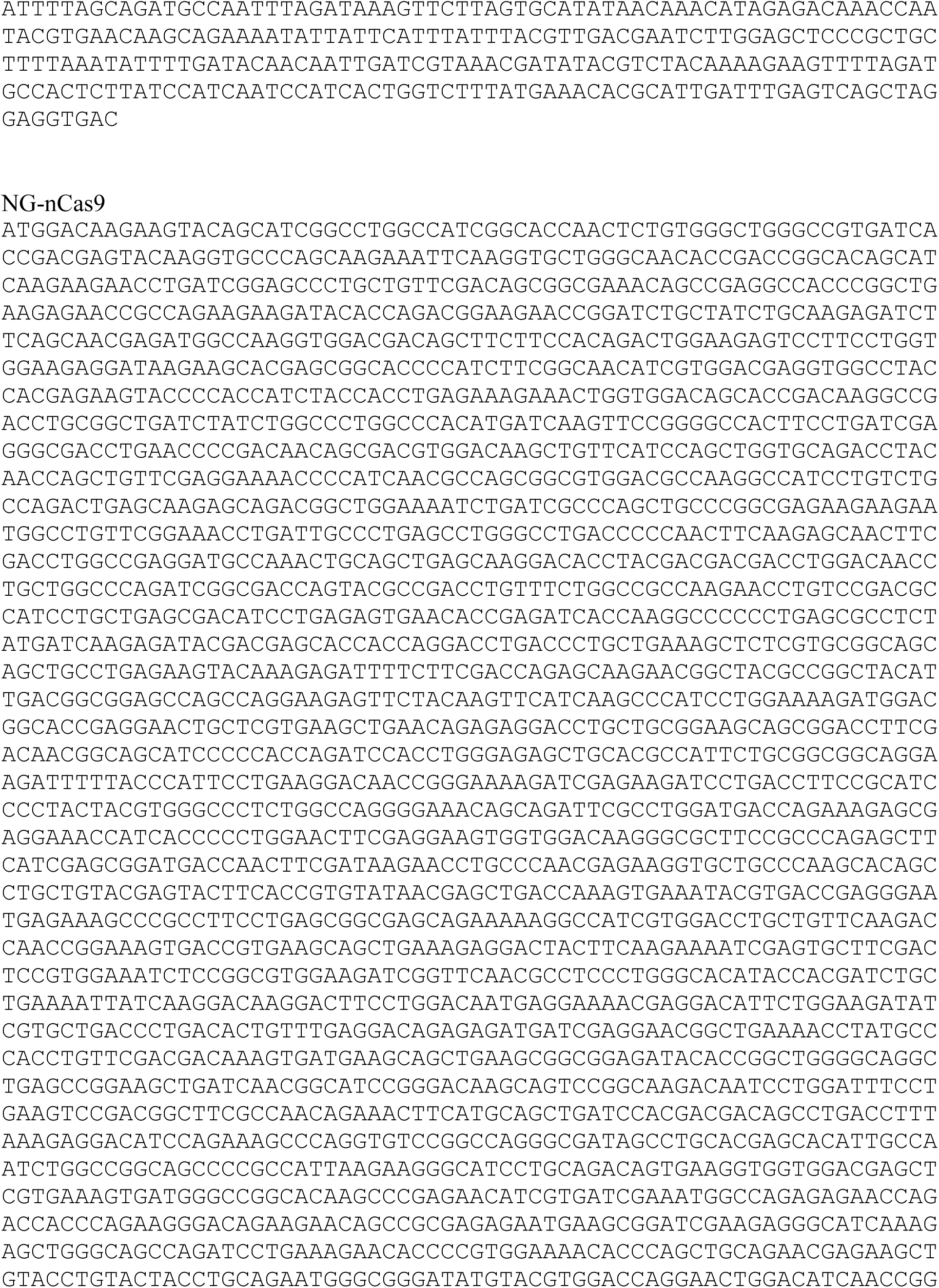
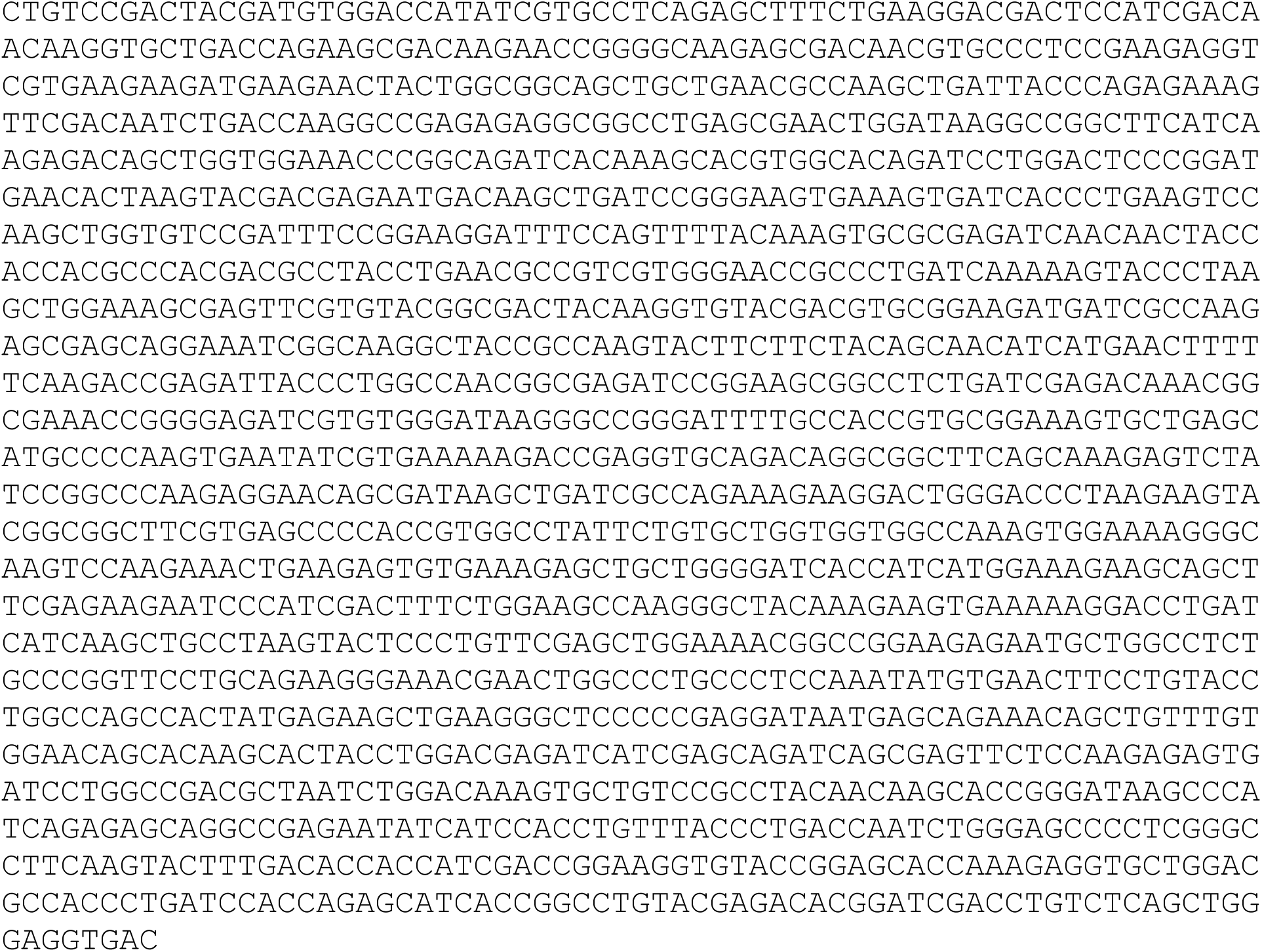

---

**Supplementary Fig 1.**
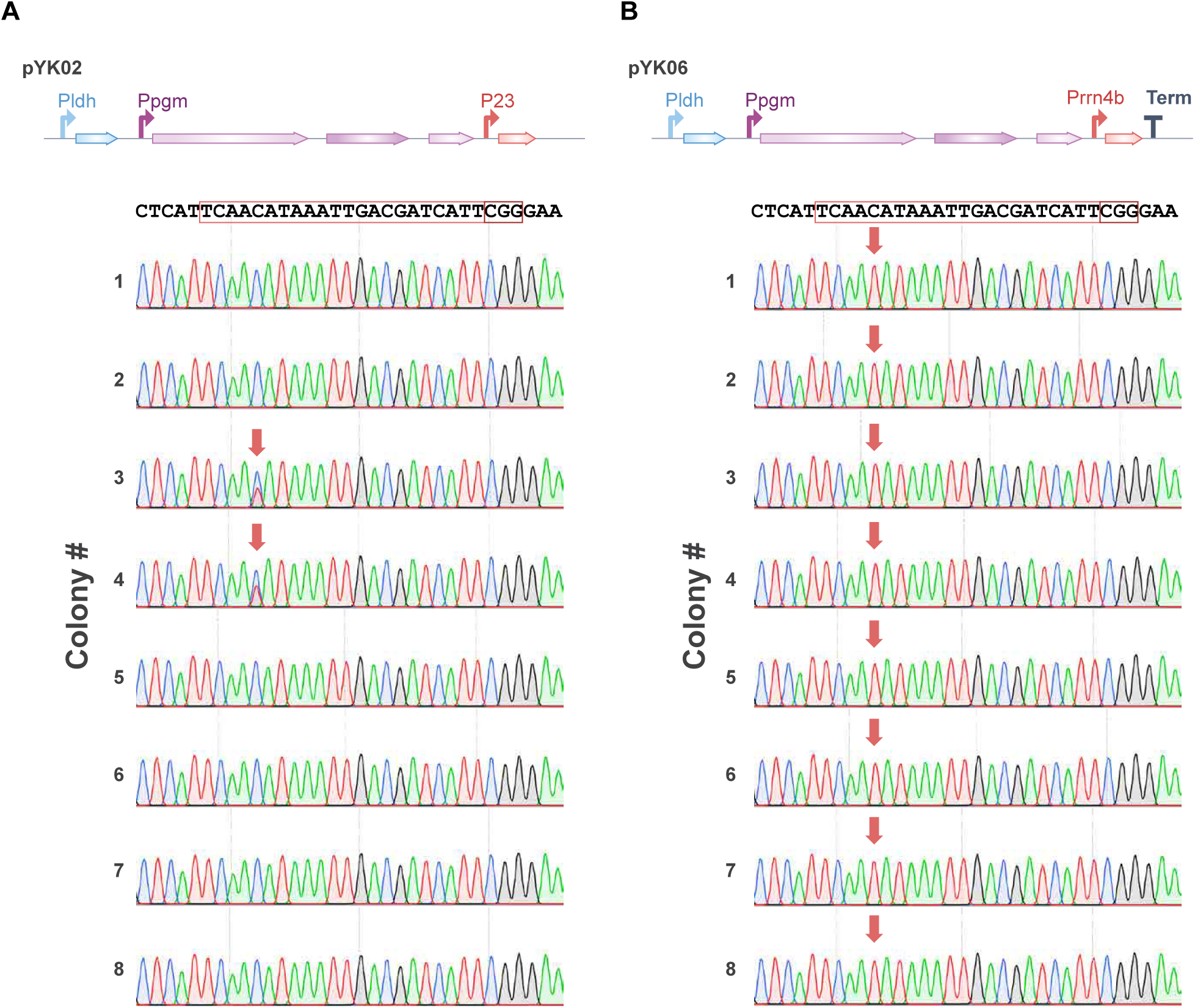
Independent sanger sequencing data for Fig. 1. Random 8 colonies of the transformants for each plasmid were sampled and the obtained sanger spectra were analyzed by EditR software to quantify mutation frequencies at each base position. Arrows indicate the detected mutations.

**Supplementary Fig 2.**
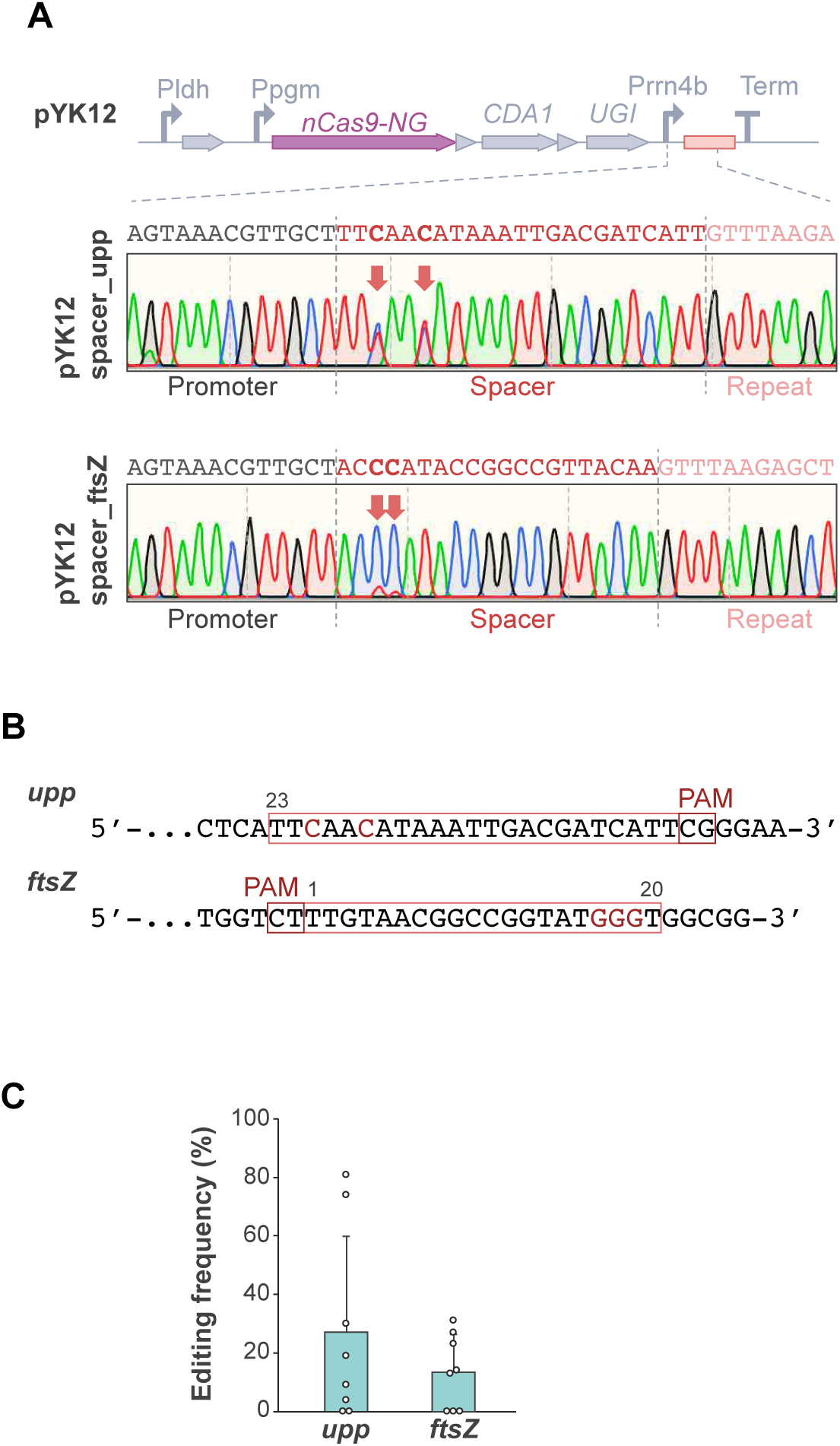
PAM-flexible Target-AID with NG-Cas9 vector and its self-targeting effect. (A) pYK12 was developed based on pYK6 by replacing coding sequence for Cas9. Sanger sequencing spectra of the constructed plasmids containing targeting sequences (*upp* (top) and *ftsZ* (bottom)) showed mixed populations at the crRNA region. (B) The target sequences for *upp* and *ftsZ* on the genome of *L. plantarum*. (C) The editing frequencies are measured by sequencing of eight randomly selected colonies and the value from the highest base positions were plotted. Averaged editing frequencies (bars) are shown with standard deviation (error bars).

**Supplementary Fig 3.**
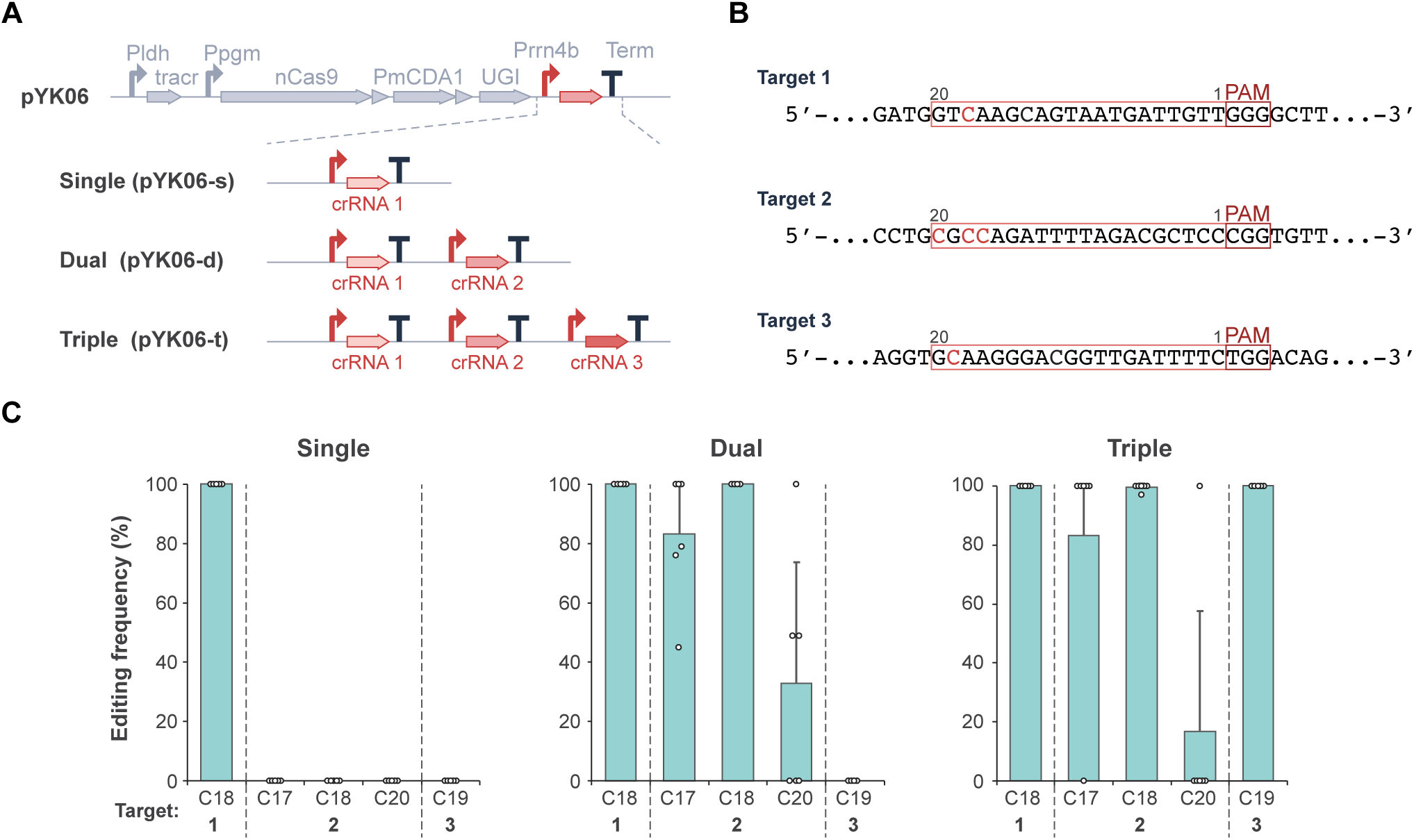
Multiplex Target-AID base editing in *L. plantarum.* (A) pYK06-s (single), -d (dual), and -t (triple) were constructed based on pYK06 by inserting in tandem each crRNA expression cassette containing a promoter, a crRNA with a target sequence and a terminator. (B) Three target sequences are selected from *urdA* gene and editable cytosines are shown in red. (C) The editing frequencies at the three sites are measured by sequencing of six randomly selected colonies and plotted at each editable base position. Averaged editing frequencies (bars) are shown with standard deviation (error bars).

**Supplementary Fig 4.**
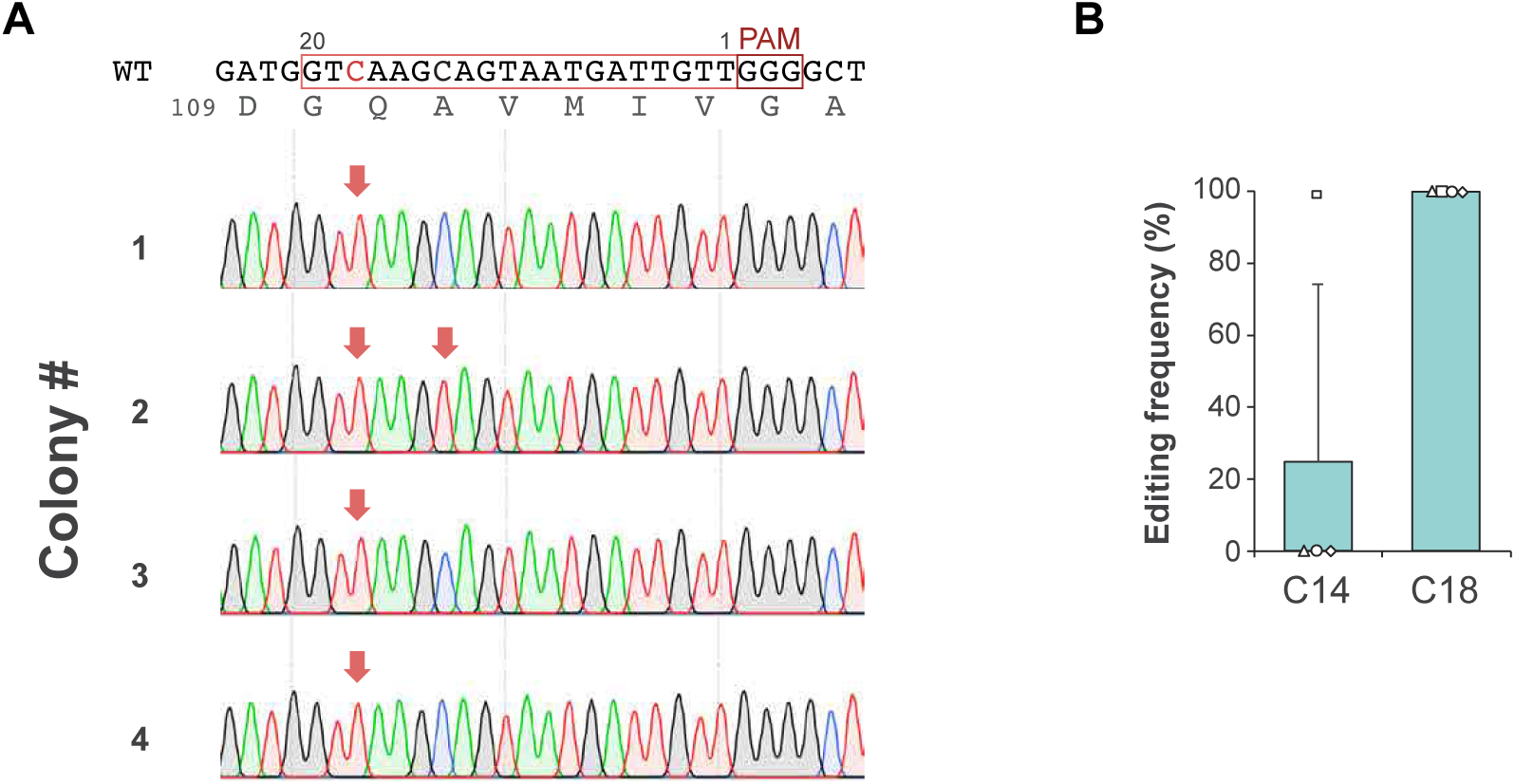
*urdA* editing with Target-AID in *L. plantarum*. (A) Sanger sequencing spectra of the four independent colonies edited at *urdA*. (B) The editing frequencies are plotted at each editable base position. Averaged editing frequencies (bars) are shown with standard deviation (error bars).

